# TEM Experiment and Complete Genome Sequencing with Comparative Genomic Analysis to Uncover Potential Probiotic Features of Isolated *Lactobacillus delbrueckii* subsp. *indicus* TY-11

**DOI:** 10.1101/2023.09.26.559573

**Authors:** Sk. Md. Jakaria Al-Mujahidy, Kirill Kryukov, Kazuho Ikeo, Kei Saito, Md. Ekhlas Uddin, Abu Ali Ibn Sina

## Abstract

**Background:** Probiotics refer to living microorganisms that exerts a variety of beneficial effects on human health. On the contrary, they also can cause infection, produce toxins within the body, and transfer antibiotic resistant genes to the other microorganisms in the digestive tract necessitating a comprehensive safety assessment. This study aimed to conduct functional genomic analysis to uncover the probiotic potential of *Lactobacillus delbrueckii* subsp. *indicus* TY-11 isolated from native yogurt in Bangladesh. We also performed Transmission Electron Microscopic (TEM) analysis, comparative genomic study as well as phylogenetic tree construction with 332 core genes from 262 genomes. These experiments on subspecies *indicus* were not studied previously.

**Results:** The strain TY-11 was identified as *Lactobacillus delbrueckii* subsp. *indicus,* whose genome (1916674bp) contained genes to adapt to diverse and stressful environments of the human gut, and no gene was identified for either antibiotic resistance or toxic metabolites. It embraced genes for the degradation of toxic metabolites, treatment of lactose intolerance, toll-like receptor 2-dependent innate immune response, heat and cold shock, bile salts tolerance in the gut, and acidic pH tolerance in the stomach. Genes were annotated for inhibiting pathogenic bacteria in the gut by inhibitory substances, (bacteriocin: Helveticin-J (331bp) and Enterolysin-A (275bp), hydrogen peroxide, and acid); blockage of adhesion sites; and competition for nutrients. Its metabolic pathway helped the human gut to recover the digestive disorders and digest indigestible nutrients. The TY-11 genome possessed additional 37 core genes of subspecies *indicus* which were deficient in the core genome of the most popular subspecies *bularicus*.

**Conclusions:** This is the first study to explore the molecular insights into intestinal residence and probiotic roles including antimicrobial activities of a representative strain (TY-11) of *Lactobacillus delbrueckii* subsp. *indicus*.

## Background

The species *Lactobacillus delbrueckii* consists at present of six subspecies, *delbrueckii, lactis, bulgaricus*, *indicus, sunki*, and *jakobsenii* showing a high level of DNA-DNA hybridization similarity but presenting a few markedly different phenotypic and genotypic characters of different ecological adaptation and restricted number of carbohydrates. Subspecies *bulgaricus, indicus,* and *lactis*, which were first isolated from dairy-based products, are all lactose-positive, whereas *delbrueckii, sunk,* and *jakobsenii* were first isolated from non-dairy-based products, are lactose negative [1]. Although *Lactobacillus delbrueckii* subsp. *bulgaricus* isolated mainly from fermented milk, it has recently also been detected in Bulgarian plants. Most isolates of the other recognized subspecies of *L. delbrueckii*, *Lactobacillus delbrueckii* subsp. *lactis*, have come from cheeses. In 2004, four strains of *Lactobacillus delbrueckii* subsp. *sunki*, designated YIT 11220, YIT 11221^T^, YIT 11466 and YIT 11673, were isolated from samples of sunki, which is a traditional, Japanese, non-salted pickle that is produced by the fermentation of the leaves of red turnips (‘otaki-kabu’) [2]. *Lactobacillus delbrueckii* subsp. *delbrueckii*, missing a lactose operon, was first observed in fermented plant extracts [3], while *Lactobacillus delbrueckii* subsp. *indicus,* which has lactose fermentation characteristic, had been isolated from fermented dairy products of India [4]. The last discovered subspecies is *Lactobacillus delbrueckii* subsp. *jakobsenii.* A novel isolate, designated ZN7a-9^T^, was named *Lactobacillus delbrueckii* subsp. *jakobsenii* and isolated from malted sorghum wort used for making an alcoholic beverage (dolo) in Burkina Faso [1]. Interestingly, while subsp. *bulgaricus, indicus* and *lactis* were all first isolated from dairy-based products, and are all lactose-positive; only two subspecies-*lactis,* and *bulgaricus* account for the commercial relevance. Subspecies *indicus* has not been marketed yet. The other three subspecies (*delbrueckii*, *sunki*, and *jakobsenii*) also have no economic pertinence, notwithstanding they are important from an evolutionary perspective because they maintain distinct nutrition and habitats. The ongoing genome sequencing of these subspecies will also aid in the illustration of further properties that probably be of future financial relevance or provide new insights on the genetic information [1].

In our study, our goal was to isolate probiotic *Lactobacillus delbrueckii* subsp. *indicus* TY-11. The subspecies *indicus* was reported previously by two different studies, where phenotypic and genotypic traits were observed by DNA–DNA hybridization, multilocus sequence typing (MLST), 16S rRNA gene sequencing, and biochemical tests [4, 5]. From the past two studies it was confirmed that subspecies *indicus* strains were potential probiotics, which fermented lactose constitutively like subspecies *bulgaricus.* However, Dellaglio *et al.* (2005) differentiated subspecies *indicus* from other five subspecies by PCR products of one or two genes, which are present in these five subspecies contrasting with subspecies *indicus.* In another point, prior studies did not focus on the responsible genes for fermentation of lactose in subspecies *indicus*. Complete genome sequence and comparative genomic study can be used to explore the mysterious genes responsible for lactose fermentation. This study is also required for the proper identification and preparing a comprehensive phylogenetic tree including all six subspecies of *Lactobacillus delbrueckii.* Moreover, this whole genome sequence is useful to explore the genes of our isolated strain for inhabitation in the human intestine, CRISPR/CRISPR-associated system (Cas), robustness in the areas of mobile genetic elements (MGEs), adaptation to environmental stress conditions, metabolism of nutrients and toxic metabolites, recovering digestive disorders, secretion system, extracellular matrix, immunity, adhesion, flavor producing capabilities, antimicrobial activities, and bacteriocin biosynthesis, which are important probiotic properties. In the previous studies, this information was not covered for the described strains of subspecies *indicus*. Functional genes and comparative genomic information from whole genome sequences of subspecies *indicus* was not studied in previous studies [4, 5]. Therefore, the aim of this study was to isolate and identify a potential strain of *L. delbrueckii* subsp. *indicus* from yogurt in Bangladesh and conduct whole genome sequencing to uncover its genomic attributes for the above-mentioned potential probiotic features. Also, Transmission Electron Microscope (TEM) was used to check the cellular morphology of the isolate.

## Results and discussion

### 1. Isolation of bacterial samples and analysis under transmission electron microscope

After morphological and 16S sequencing result analysis, strain TY-11 was selected for further studies. Rod shaped cells of varying lengths and widths (2.2 µm −9µm lengths and 0.7 µm −1.3µm widths) with rounded ends were observed under TEM. Two cells of TY-11 were adhered by exopolysaccharide like ingredients as shown in the figure **(Fig: 1)**. It is the first TEM analysis report for a strain of subsp. *indicus*.

**Fig: 1.**
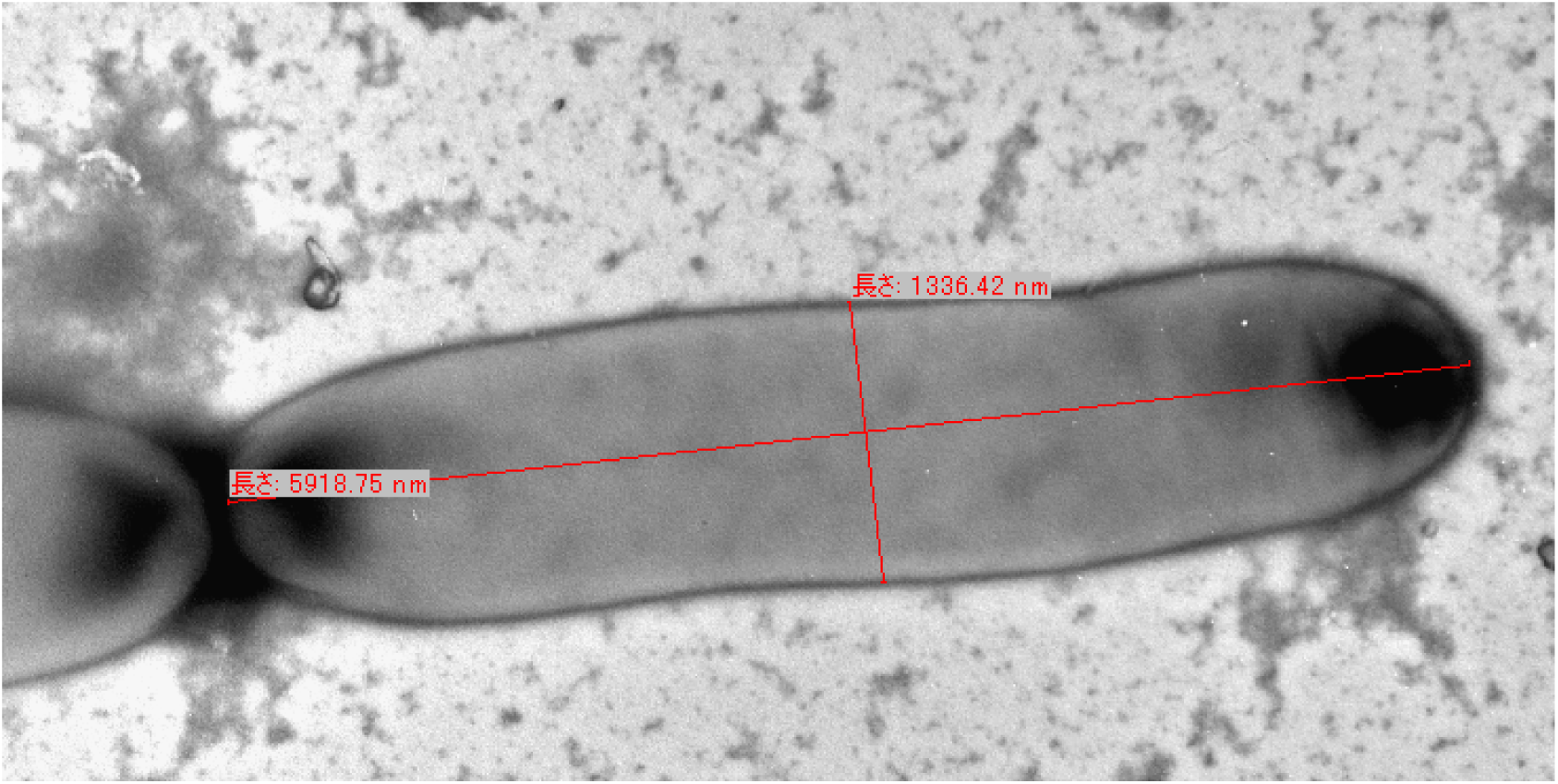
*L. delbrueckii* subsp. *indicus* strain TY-11 at 20,000 magnifications displayed cell length=5979.89nm and width= 1337.42nm.

### 2. QC check of genome sequence

According to the FastQC report, the total number of reads is 4135806 (2067903 pairs), and their total size was 1.3 GB. Sequence length was 35-53, and %GC was 49. According to the report of fastp (version 0.19.7), mean length before filtering was 134bp, 135bp and after filtering it was 131bp, 131bp. Duplication rate was 0.944674%. Insert size was 35.

### 3. Genome composition analysis and genome visualization by assembly and annotation

In the TY-11 genome, the total length of coding regions had 1916674 bases. The total length of coding regions accounted for 99.48% of the whole genome, whereas noncoding regions were comprised of only 0.52% nucleotides of the whole genome. The genome had 122 contigs, 69 tRNA, 4 rRNA, 1 tmRNA, and 1911 CDS including 74 hypothetical genes, and 84 pseudo genes **(Fig: 2, Additional file 1: Table S1)**. The TY-11 genome contained a total forty-five CRISPR/CRISPR-associated system (Cas) genes. It had a total 105 mobile elements. Antibiotic resistance gene was not identified in the TY-11 genome.

**Fig: 2.**
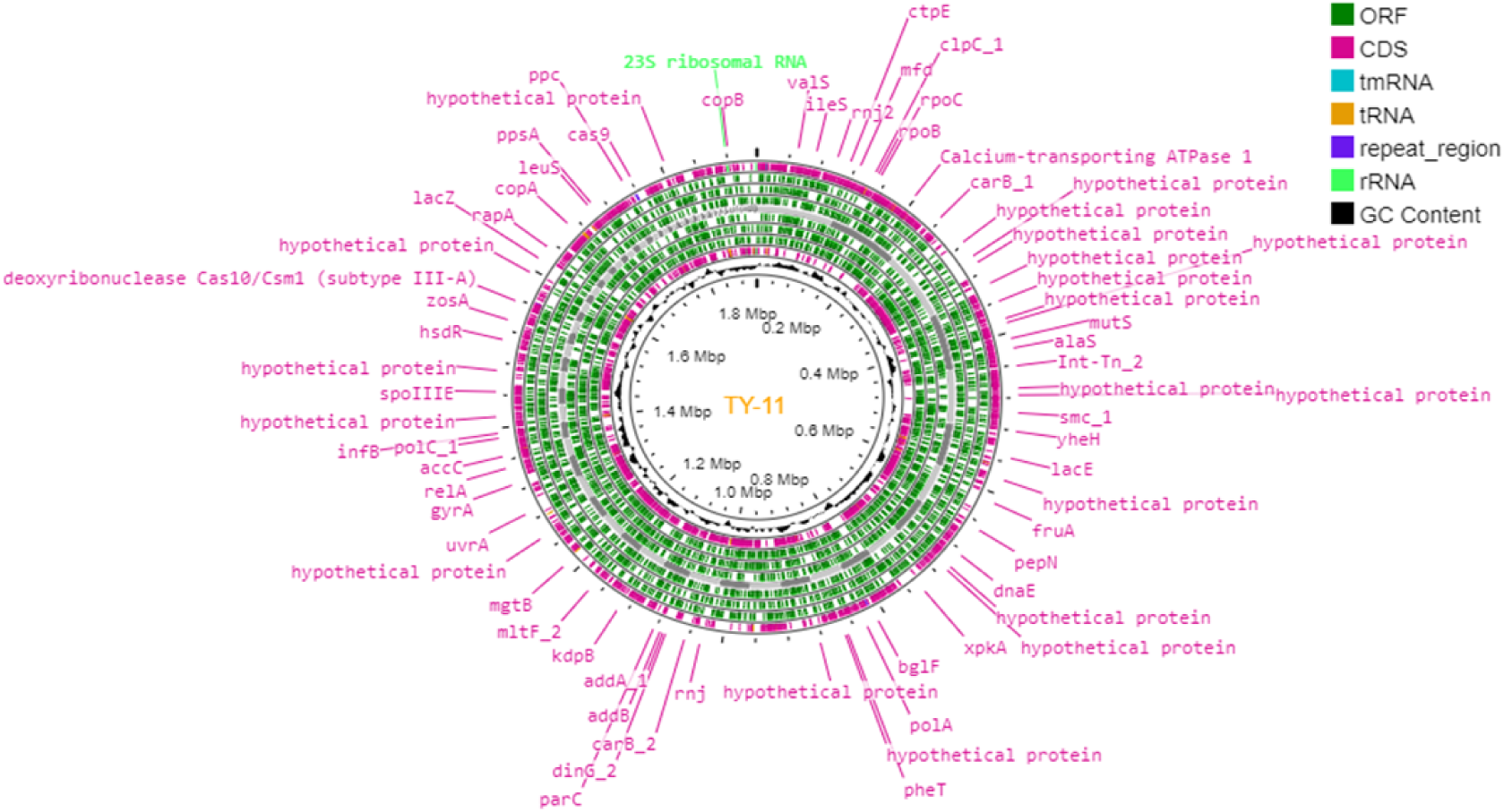
Circular genome map of TY-11 genome. Genome Visualization ID: 3fe48947-6842-4b90-8fe1-9020448e1541. The circular map was generated using CGViewBuilder (version 1.1.1). The information marked in the figure, is depicted from the external band to inmost as follows: CDSs (forward strand), ORF (forward strand), genome size, ORF (reverse strand), CDSs (reverse strand), GC (guanine–cytosine) content. The CRISPR/Cas systems are assorted into two classes according to the structure and function of Cas protein: class I and class II. They are further split into six subclasses (type I– VI). Type I, III, and IV are involved in Class I, and type II, V, and VI are included in class II [6]. The genome had twenty-seven Cas genes, and eleven Cas clusters. **(Additional file 2: Table S2)**. The TY-11 genome contained seven CRISPR arrays **(Additional file 2: Table S2, Fig: 3)**. The genome stability of a strain and, thereby, its adjustment in the environment is improved by the CRISPR loci [7].

**Fig: 3.**
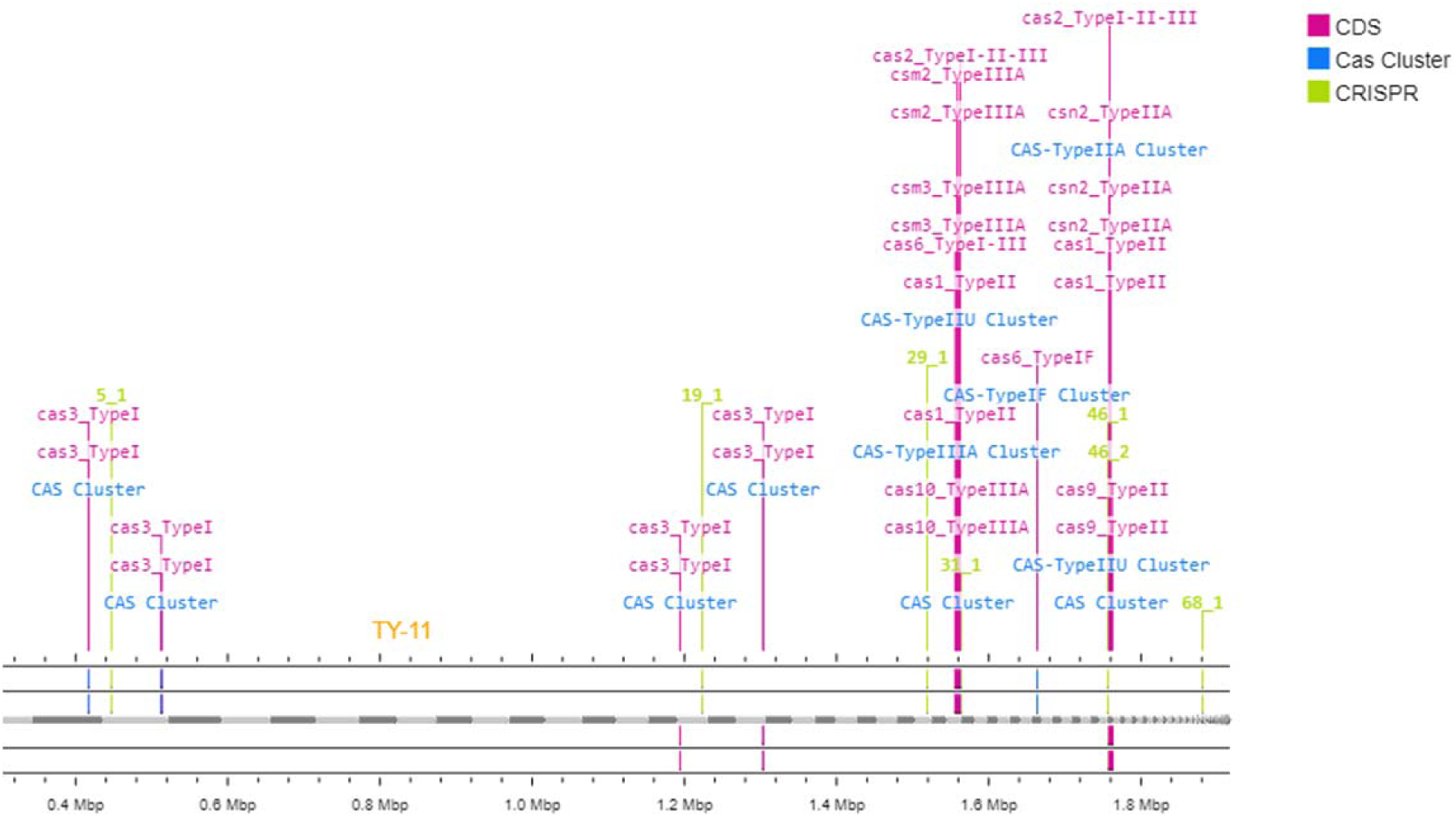
CRISPR/CRISPR-associated system (Cas) genes. The TY-11 genome contained seven CRISPR arrays (green), twenty-seven Cas genes (dark pink), and eleven Cas clusters (blue).

### 4. CRISPR/CRISPR-associated system (Cas) genes

In the TY-11 genome, the tool detected total forty-five CRISPR/CRISPR-associated system (Cas) genes, which were of type I, II, and III (cas3_TypeI, cas10_TypeIIIA, csm2_TypeIIIA, csm3_TypeIIIA, cas2_TypeI-II-III, cas6_TypeI-III, cas1_TypeII, cas6_TypeIF, cas2_TypeI-II-III, cas9_TypeII, csn2_TypeIIA, csn2_TypeIIA, and csn2_TypeIIA).

### 5. Mobile genetic elements

To facilitate mobile genetic elements (MGEs) annotation, mobile orthologous group database (mobileOG-db) was developed. mobileOG-db is available at mobileogdb.flsi.cloud.vt.edu/, where researcher can inspect the dynamic mobilome subsets [8]. There was no complete prophage found in TY-11 genome. Twenty genes encoded by phage were predicted as Major mobileOG Category in the genome (**Additional file 3: Table S3**). Among them, three genes were hypothetical genes having nucleotide lengths of 152bp, 313bp, and 154bp. Other seventeen genes were identified as clpX (418bp), clpP (195bp), clpC (732bp), clpB (697bp), thyA (319bp), ftsH (738bp), mur (294bp), orf36 (479bp), orf35 (405bp), orf623 (624bp), erf (232bp), groS (95bp), groL (538bp), lysA (298bp), lexA (207bp), dnaK (615bp), and nusA (407bp) **(Fig: 4)**.

**Fig: 4.**
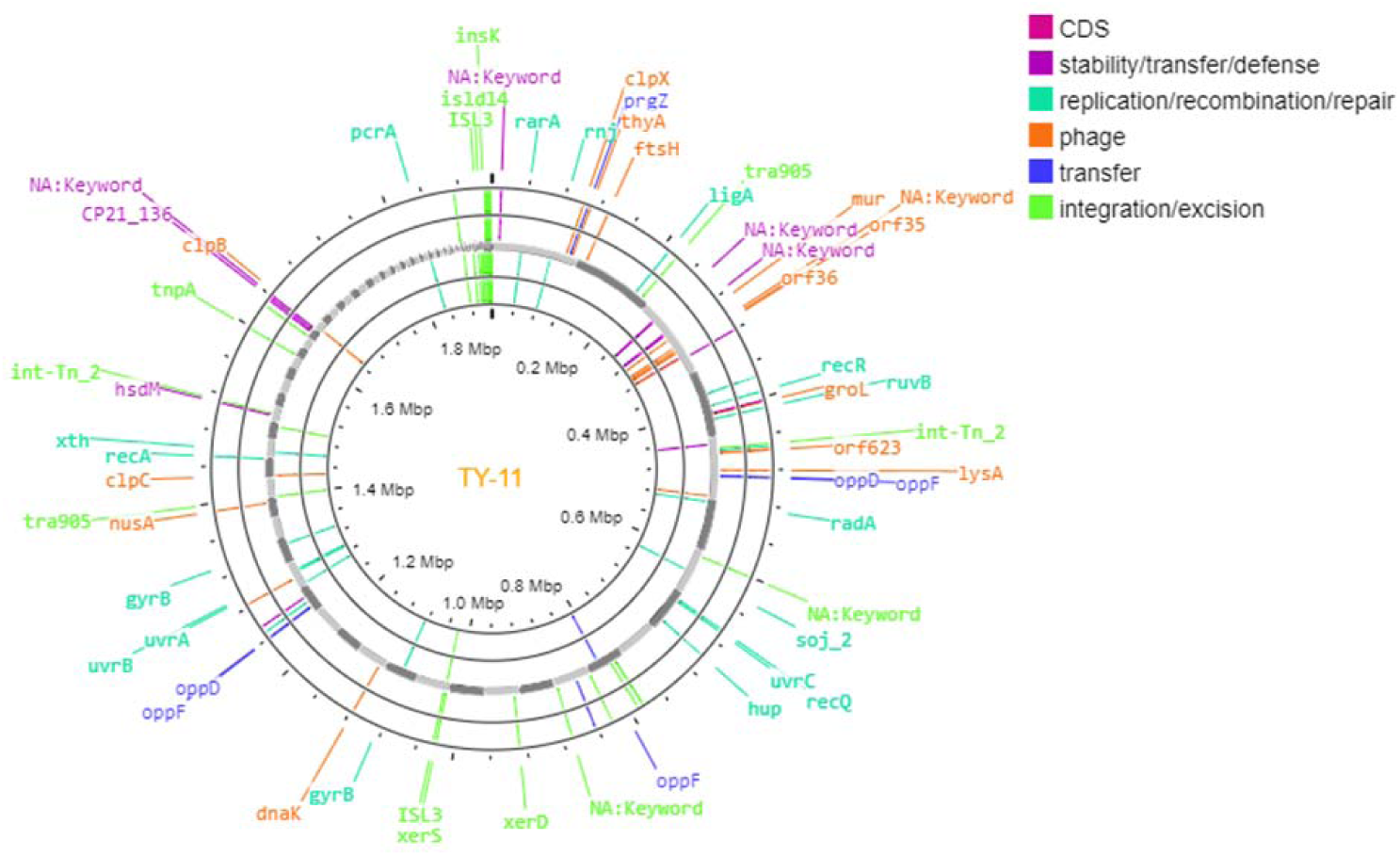
Mobile genetic elements in the genome TY-11. The map was formed in circular shape using CGViewBuilder (version 1.1.1). The labelled information is presented as follows: repeat region, replication/recombination/repair, phage, transfer, integration/excision, stability/transfer/defence, and ORF.

### 6. Phylogenetic study and identification

Phylogenetic study was conducted for finding the evolutionary relationships of taxa. The new genome TY-11 was at the bottom in the tree, and the most closely related 3 unspecified subspecies genomes are mentioned as follows:

GCF_025186005.1 Lactobacillus delbrueckii = Lactobacillus delbrueckii strain 862, GCF_025193525.1 Lactobacillus delbrueckii = Lactobacillus delbrueckii strain CIRM-BIA 865, and GCF_021600505.1 Lactobacillus delbrueckii = Lactobacillus delbrueckii strain ME-792.

The next nearest were 3 genomes of L. delbrueckii subsp. indicus. These 7 genomes were organized in a cluster with a rather long branch to the rest of the tree. It was assumed that all 7 are probably subsp. *indicus*. In the first tree, there were total 4 *Lactobacillus delbrueckii* subsp. *indicus*, except TY-11 genome. The nearest three sequences of *Lactobacillus delbrueckii* subsp. *indicus* (GCF_001908415.1= JCM 15610, GCF_001435795.1= DSM 15996, GCF_001189855.1 =JCM 15610) stayed in a cluster. GCF_025960345.1= S2 was placed outside of this cluster. This could be a misclassification **(Additional file 4: Fig S1).** In the second tree, it showed same feature, that is, GCF_025960345.1= S2 was in the out-group, and other three subsp. *indicus* genomes stayed in one cluster near to TY-11 **(Fig: 5)**. Overall, from the tree of 262 genomes, it seems that TY11 belongs to *Lactobacillus delbrueckii* subsp. *indicus*.

**Fig: 5.**
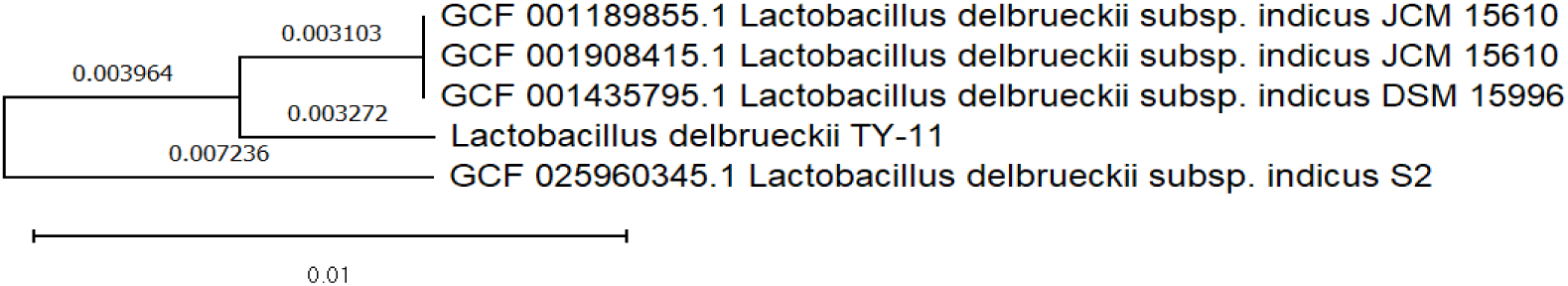
Phylogenetic tree-2. The tree was inferred using the Neighbor-Joining method employing 5 genomes of *L. delbrueckii* subsp. *indicus* (based on aligned amino-acid sequences of 353 common genes of TY-11 genome and 4 subsp. *indicus* genomes).

DNA-DNA hybridization (DDH) is used for the delineation of bacterial strains. Conspecific genomes must show >70% DDH similarity following the rules of bacterial nomenclature code. Progressively, DDH is accompanied with or substituted by average nucleotide identity (ANI) comparisons (about >70% DDH similarity corresponds to >94% of ANI in the core genome and >96% in universal marker genes). Within-species ANI values are larger than 97% in the most cases [9]. The complete genome of TY-11 showed 98.980% ANI with that of the closest microorganism, *Lactobacillus delbrueckii* subsp. *indicus* JCM 15610 (GCA_001908415.1=GCF_001908415.1= ASM190841v1) (RAPT job ID: 669b682-urs-6d54), whereas *L. delbrueckii* subsp. *jakobsenii* ZN7a-9 = DSM 26046 was a strain of the farthest subspecies having 97.404% ANI.

The strains of six subspecies, *delbrueckii, lactis, bulgaricus*, *indicus, sunki*, and *jakobsenii* represented a little variation in the ANI percentage (**Table: 1)**. The core genome sequence (353genes) of TY-11 showed 99.01441% similarity with the cluster of all three nearest sequences of *L. delbrueckii* subsp. *indicus* (GCF_001908415.1= JCM 15610, GCF_001435795.1= DSM 15996, GCF_001189855.1 =JCM 15610), while it showed 98.32962% similarity with the second nearest *L. delbrueckii* subsp. *Indicus* (GCF_025960345.1= S2). Therefore, TY-11 was identified as *L. delbrueckii* subsp. *indicus*.

**Table: 1.** Average nucleotide identity of the strain TY-11 with the nearest strains of six subspecies in the species *L. delbrueckii*.

| Organism name | Subspecies name | GenBank assembly accession | Query percentage coverage | ANI |
| --- | --- | --- | --- | --- |
| *L. delbrueckii* subsp. *indicus* JCM 15610 | indicus | GCA_001908415.1 | 85.1 | 98.98 |
| *L. delbrueckii* subsp. *lactis* DSM 20072 | lactis | GCA_002278095.1 | 81.3 | 97.338 |
| *L. delbrueckii* subsp. *sunkii* | sunkii | GCA_001888965.1 | 80.4 | 97.519 |
| *L. delbrueckii* subsp. *delbrueckii* | delbrueckii | GCA_006740305.1 | 79.5 | 97.39 |
| *L. delbrueckii* subsp. *bulgaricus* ATCC 11842 = JCM 1002 | bulgaricus | GCA_000056065.1 | 78.2 | 97.241 |
| *L. delbrueckii* subsp. *jakobsenii* ZN7a-9 = DSM 26046 | jakobsenii | GCA_001888925.1 | 77.0 | 97.404 |

**Table: 2.** Central carbohydrate metabolism.

| Name of pathway | Prokka ID | KEGG orthologs | Gene name |
| --- | --- | --- | --- |
| Glycolysis (Embden-Meyerhof pathway)(M00001) | LPJCKDFM_01631 | K25026 | Glucokinase |
|  | LPJCKDFM_00021 | K01810 | glucose-6-phosphate isomerase |
|  | LPJCKDFM_00746 | K00850 | 6-phosphofructokinase 1 |
|  | LPJCKDFM_00629 | K21071 | ATP-dependent phosphofructokinase / diphosphate-dependent phosphofructokinase |
|  | LPJCKDFM_01453 | K01624 | fructose-bisphosphate aldolase, class II |
|  | LPJCKDFM_01320 | K01803 | triosephosphate isomerase (TIM) |
|  | LPJCKDFM_01322 | K00134 | glyceraldehyde 3-phosphate dehydrogenase (phosphorylating) |
|  | LPJCKDFM_01321 | K00927 | phosphoglycerate kinase |
|  | LPJCKDFM_00582, LPJCKDFM_00945 | K01834 | 2,3-bisphosphoglycerate-dependent phosphoglycerate mutase |
|  | LPJCKDFM_01182 | K01689 | Enolase |
| Pentose phosphate pathway, oxidative phase, glucose 6P => ribulose 5P (M00006) | LPJCKDFM_01842 | K00033 | 6-phosphogluconate dehydrogenase |
|  | LPJCKDFM_00456 | K00036 | glucose-6-phosphate 1-dehydrogenase |
|  | LPJCKDFM_01488 | K07404 | 6-phosphogluconolactonase |
| The three-carbon compound core module of glycolysis (M00002) | LPJCKDFM_01322 | K00134 | (phosphorylating) the enzyme glyceraldehyde 3-phosphate dehydrogenase |
|  | LPJCKDFM_00747 | K00873 | pyruvate kinase |
|  | LPJCKDFM_01321 | K00927 | phosphoglycerate kinase |
|  | LPJCKDFM_01182 | K01689 | Enolase |
|  | LPJCKDFM_01320 | K01803 | triosephosphate isomerase |
|  | LPJCKDFM_00582, LPJCKDFM_00945 | K01834 | 2,3-bisphosphoglycerate-dependent phosphoglycerate mutase |

### 7. Functionality analysis of genome by KEGG annotations

According to KEGG Mapper Reconstruction Result, total 1079 genes (56.5%) were annotated from 1911 CDS **(Fig: 6)**. The genes (56.5%), annotated by KEGG annotations, were cited in this study as KEGG ID. The remaining genes (43.5) from PROKKA (version 1.13) annotation were cited in this study mentioning PROKKA ID [10].

**Fig: 6.**
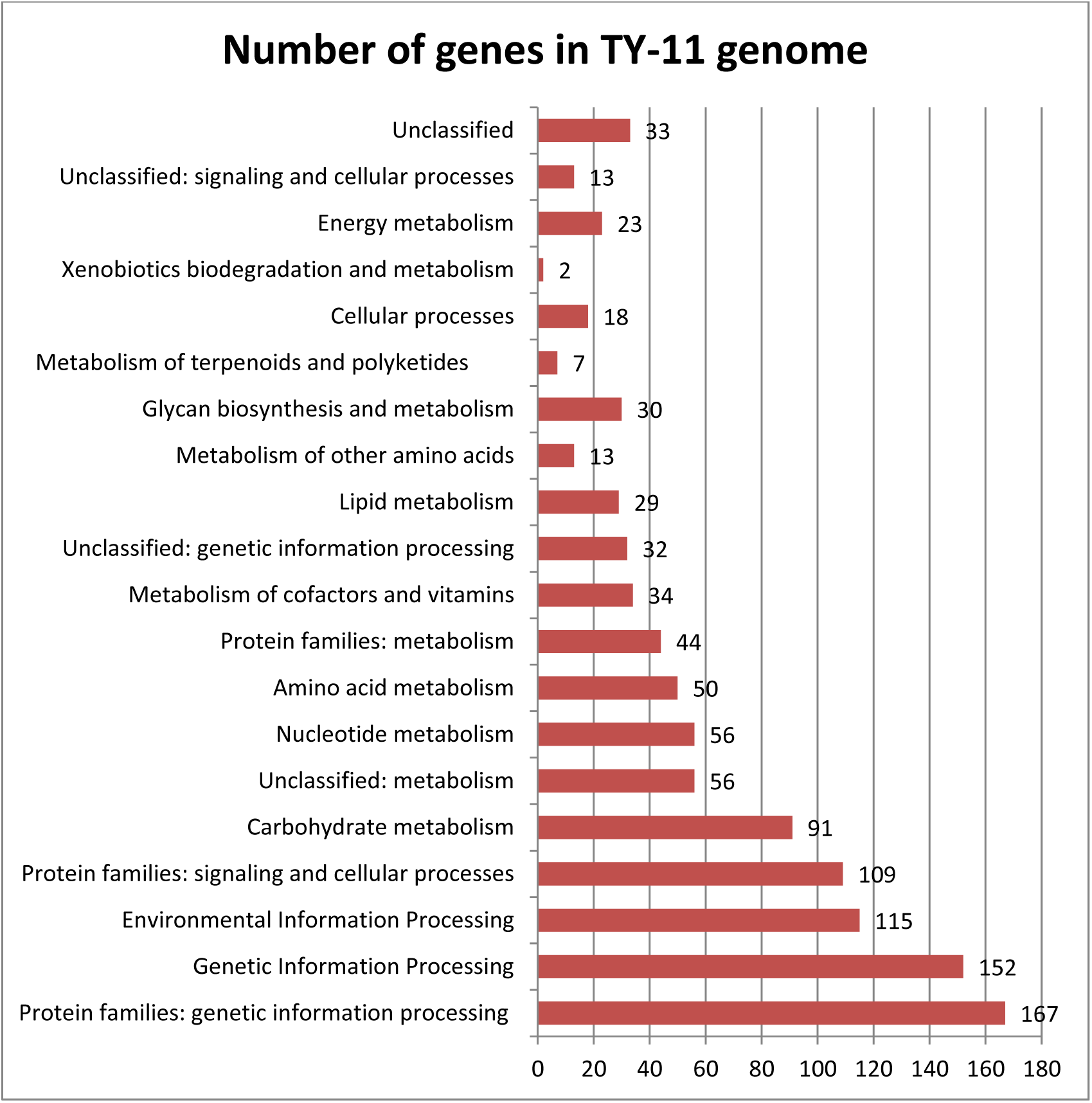
Functionality analysis of TY-11 genome by KEGG annotations. Number of genes by KEGG annotations of TY-11 genome has been showed in the figure.

#### 7.1.1. Central carbohydrate metabolism

TY11 embraced a complete set of genes (11genes) of Embden-Meyerhof pathway for glycolysis to convert glucose to pyruvate (M00001). It had a complete set of genes (6 genes) for glycolysis core module involving three-carbon compounds (M00002). It gained Pentose phosphate pathway, oxidative phase, to convert glucose 6P to ribulose.

#### 7.1.2. Anaerobic growth

TY-11 was an anaerobic bacterium according to genomic data, because it had lactose dehydrogenase gene, L-lactate dehydrogenase (K00016) by which it is able to change pyruvate into l-lactate. One of the H transfer (oxidoreductase) enzymes, lactate dehydrogenase uses NADH to catalyze the reversible conversion of pyruvate to lactate. The enzyme essentially participates in the anaerobic metabolism of glucose when oxygen is absent or scarce [11].

Pyruvate + NADH + H+ --> Lactate + NAD+

The anaerobic ribonucleoside-triphosphate reductase activating protein (LPJCKDFM_00962) found in TY-11 is activated under anaerobic conditions by the production of an organic free radical, which uses reduced flavodoxin and S-adenosylmethionine as cosubstrates to generate 5’-deoxy-adenosine [12]. It also had anaerobic ribonucleoside-triphosphate reductase (LPJCKDFM_00963=K21636), which has an oxygen-sensitive activity that reduces CTP to dCTP in the presence of NADPH, dithiothreitol, Mg2+ ions, and ATP [13]. The anaerobic character of TY-11 is consistent with the anaerobic environment in the human gut, because 99.9% of colonic microflora are obligate anaerobes [14].

#### 7.1.3. Lactose metabolism and lactose intolerance

According to the present study, TY-11 had lactose transporting PTS system (K02786 and K02788) (discussion in Section 7.8), which simultaneously transports lactose from the periplasm or extracellular space into the cytoplasm and phosphorylates it. TY-11 also had 6-phospho-beta-galactosidase (lacG=K01220), which produced glucose and galactose-6-phosphate (Gal-6P) during the hydrolysis of lactose-6-phosphate (Lac-6P) [15, 16]. This PTS system, the phospho-β-galactosidase and the galactose-6-P metabolizing enzymes appear to be rare in the lactobacilli of the acidophilus group and in lactobacilli in general.

In the TY-11 genome, lactose permease enzyme was missing, although it retained lactose utilizing protein: glycoside hydrolase family 1 protein (LPJCKDFM_01304=K01223), glycoside hydrolase family 68 protein, glycoside hydrolase (LPJCKDFM_00424), and glycoside hydrolase family 73 protein (LPJCKDFM_00227), which are the enzymes with a number of known activities including beta (EC 3.2.1.23) galactosidase [17].

#### 7.2. Adaptation to heat-shock and cold-shock stress

TY-11 beard various genes encoding heat stress proteins, for example, Hsp33 family molecular chaperone HslO (LPJCKDFM_00136), molecular chaperone DnaJ (LPJCKDFM_01166), molecular chaperone DnaK (K04043), chaperonin GroEL (K04077), and co-chaperone GroES (LPJCKDFM_00449) [7]. GroEL is a protein which is required for the proper folding of many proteins at different temperature conditions (Section 5). In *Mycobacterium smegmatis*, it was shown that DnaK has a non-redundant role in the folding of essential proteins and in generally maintaining protein homeostasis at both high and low temperatures and is therefore required even in the absence of stress [18]. Like *M*. *smegmatis*, the α-proteobacterium *Caulobacter crescentus*, which is a model organism for studying bacterial cell cycle regulation, strictly depends on DnaK at all temperature conditions [19]. This genome had groS (95bp), and groL (538bp) genes (described in Section-4), and a gene (LPJCKDFM_01163) for heat-shock repressor protein HrcA. It had a defence mechanism against a sudden heat-shock stress [20]. It had cold shock domain-containing protein (LPJCKDFM_00078). These so-called ‘cold shock’ proteins are thought to help the cell to survive in temperatures lower than optimum growth temperature, by contrast with heat shock proteins, which help the cell to survive in temperatures greater than the optimum [21].

#### 7.3. Tolerance to weak acids and bile salts in the gut

The TY-11 genome had choloylglycine hydrolase (K01442). These proteins are associated with tolerance to acids and bile salts in the gut. Certain species of the indigenous microflora, including a number of lactobacilli and bifidobacteria, have evolved the ability to deconjugate bile salts by bile salt hydrolase (BSH) [22]. However, The TY-11 genome lacked this enzyme.

#### 7.4. pH homeostasis under acid stress

TY-11 had F0F1 ATP synthase subunit A, B, C, alpha, beta, gamma, delta, and epsilon (LPJCKDFM_00033-40). Low driving force causes ATP synthases to act as ATPases and produce a transmembrane ion gradient at the expense of ATP hydrolysis [23]. This genome had chaperonin GroEL (K04077) and co-chaperone GroES (LPJCKDFM_00449) (discussion in Section 5). These general stress proteins, are overexpressed during acid stress [24].

#### 7.5. Lifestyle adaptation to other stress

The TY-11 genome may contribute to the tolerance of oxidative stress through the coding of the protein thioredoxin-disulfide reductase (LPJCKDFM_01338=K00384) and glutaredoxin (LPJCKDFM_00958) that catalyse glutathione-dependent disulphide reductions. The TY-11 genome contains genes that code for stress-related proteases like the ATP-dependent proteases clpX (418 bp) and clpP (195 bp) (discussed in Section 5) and subunit HslV (LPJCKDFM_00980), which prevent abnormal protein damage [7]. It had genes for osmoprotectant transport system permease protein (LPJCKDFM_00370, LPJCKDFM_00931, and LPJCKDFM_00933=K05846) which play crucial roles in the adaptation of cells to various adverse environmental conditions [25]. Peroxide stress protein YaaA (LPJCKDFM_00708) was predicted to involve in the cellular response to hydrogen peroxide (H_2_O_2_) stress. By reducing the amount of unincorporated iron in the cell, YaaA protects DNA and proteins from oxidative damage caused by H_2_O_2_ stress [12].

#### 7.6. Biosynthesis system **of** amino acid

Complete threonine pathway, employing 5 genes (M00018), was predicted in TY11 genome: aspartate → homoserine →threonine. It had the gene sets to complete the pathway for proline metabolism (3 genes for M00015) (**Table: 3**).

**Table: 3.** Threonine and proline metabolism.

| Name of pathway | Prokka ID | KEGG orthologs | Gene name | Gene symbol |
| --- | --- | --- | --- | --- |
| (M00018) The synthesis of threonine, aspartate → homoserine → threonine | LPJCKDFM_01420 | K00003 | homoserine dehydrogenase<br>[EC:1.1.1.3] | hom |
|  | LPJCKDFM_01422 | K00133 | aspartate-semialdehyde dehydrogenase<br>[EC:1.2.1.11] | asd |
|  | LPJCKDFM_00621 | K00872 | homoserine kinase<br>[EC:2.7.1.39] | thrB |
|  | LPJCKDFM_00284, LPJCKDFM_01421 | K00928 | aspartate kinase<br>[EC:2.7.2.4] | lysC |
|  | LPJCKDFM_00622 | K01733 | threonine synthase<br>[EC:4.2.3.1] | thrC |
| (M00015) Proline biosynthesis, glutamate ⇒ proline | LPJCKDFM_00742 | K00147 | glutamate-5-semialdehyde dehydrogenase<br>[EC:1.2.1.41] | proA |
|  | LPJCKDFM_00741 | K00286 | pyrroline-5-carboxylate reductase<br>[EC:1.5.1.2] | proC |
|  | LPJCKDFM_00743 | K00931 | glutamate 5-kinase | proB |

This probiotic had twelve genes to complete the entire acetyl-DAP pathway for lysine metabolism from aspartate (Ko00300, M00525): K00928, K00133, K01714, K00215, K05822, K00841, K05823, K01778, K01586, and K05825 (**Fig: 7**). Lysine is an essential amino acid for human [26]. It had the complete gene sets for arginine metabolism (Ko00220, M00844): K00611, K01940, and K01755(**Fig: 8**). Arginine is a conditionally essential amino acid for human [27]. It possessed a better transport system to make up for its restricted ability to synthesize amino acids [7].

**Fig: 7.**
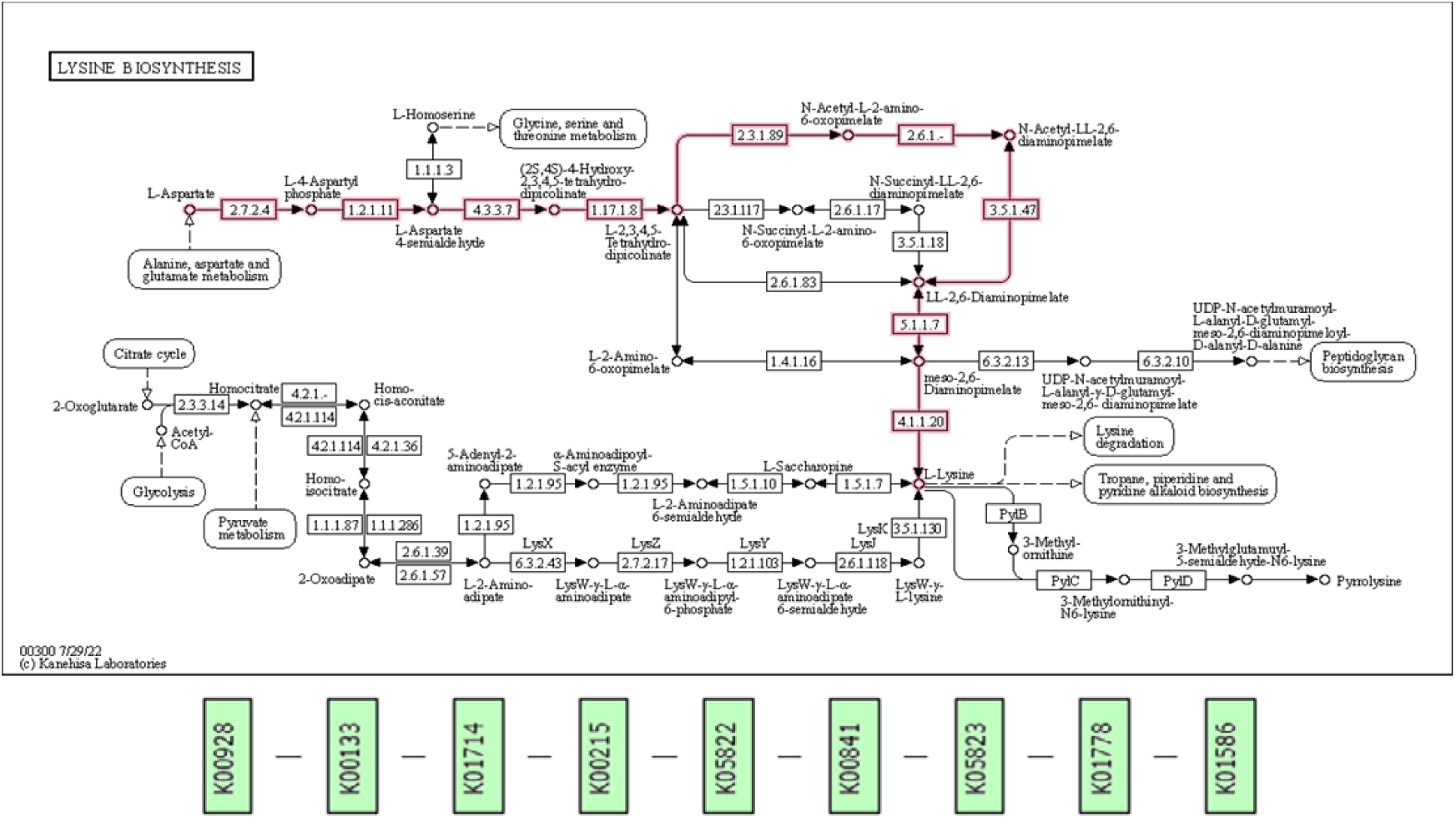
Lysine biosynthesis. The upper flowchart shows lysine biosynthesis pathway of TY-11 genome in purple colour. The KEGG orthologs of the enzymes involved in lysine biosynthesis pathway (upper flowchart) are represented serially from the left to the right in the lower flowchart.

**Fig: 8.**
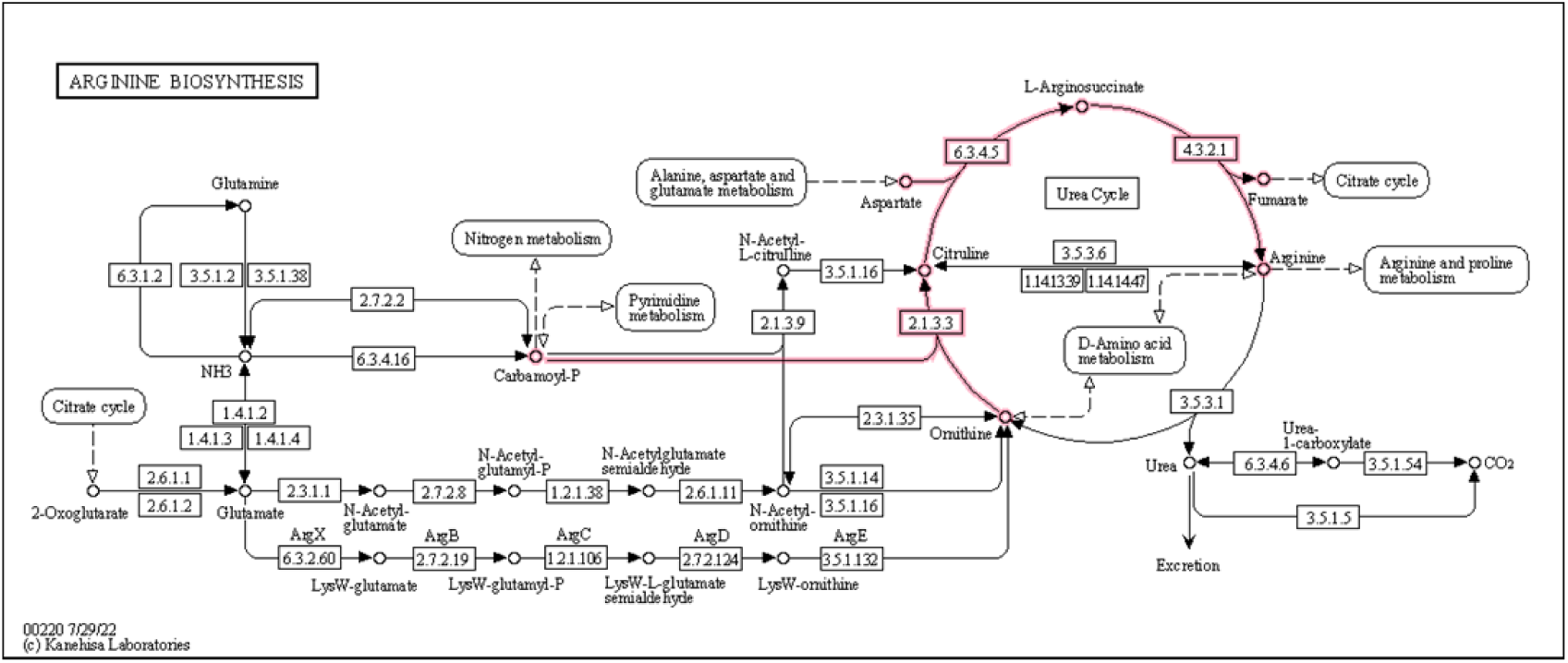
Arginine biosynthesis. The diagram depicts arginine biosynthesis pathway of TY-11 genome in purple colour. The enzymes involved in arginine biosynthesis are as follows: 21.33=K00611, 63.45=K01940, and 43.21=K01755.

#### 7.7. Metabolism of cofactors and vitamins

TY-11 genome had four genes encoding enzymes for Coenzyme A biosynthesis (M00120) from pantothenate. All genes for folate biosynthesis pathway were annotated in TY-11 genome (Ko:00790) to synthesize folate from Guanosine-5’-triphosphate (GTP) except alkaline phosphatase (EC: 3.1.3.1). TY-11 genome entailed dihydrofolate reductase (K00287), which converts folate to dihydrofolate, and then dihydrofolate to tetrahydrofolate (THF) **(Additional file 6: Table S5**). In cells, folic acid (vitamin B9) is reduced to THF, a biologically active form [28]. The folate deficiency manifests in fatigue, muscle weakness, tingling of extremities, and loss of joint position/coordination in human [29, 30].

#### 7.8. Transport system

The ABC transporter system and PTS were represented by 78 genes in the TY-11 genome, which was around 1.9 Mb in size. The genome harboured 62 genes that coded for the elements of the ABC transporter system (**Additional file 5: Table S4**). The number of transporters in a genome is generally proportional to genome size [31]. ABC transporter proteins are by far the most abundant of transporters, typically accounting for half of the transporters in a bacterial genome. The 4.6 Mb genome of *Escherichia coli* encodes 78 ABC transporter systems, which is typical for a genome of this size. *Agrobacterium tumefaciens* has a 5.7 Mb genome that encodes over 200 ABC systems, in contrast to *Mycobacterium tuberculosis’*s 4.4 Mb genome that only encodes 38 ABC systems. [31].

Many of these TY-11genome ABC importers transport inorganic ions, amino acids, and peptides; however, it is unknown what kinds of substances the majority of the exporters export. The genome contained 11 amino acid-permease type transporters and one transporter for branched-chain amino acids, which is the typical number of transporters found in the majority of other lactic acid bacteria [7]. The genes encoding the flippase proteins (LPJCKDFM_01660 flippase-like domain-containing protein, LPJCKDFM_01863&00849 flippase, and LPJCKDFM_00212 oligosaccharide flippase family protein) were also discovered in the TY-11 genome. Flippases, which are infrequently written flipases, are ABC transporter or P4-type ATPase families of transmembrane lipid transporter proteins that are found in membranes. They help phospholipid molecules flow across the two leaflets that make up a cell’s membrane [32].

The strain TY-11 had osmoprotectant transport system permease protein (K05846). Osmoprotectants (also termed compatible solutes) are highly soluble compounds that carry no net charge at physiological pH and are nontoxic at high concentrations. When salt concentrations or temperatures are unfavourable, osmoprotectants help to increase osmotic pressure in the cytoplasm and can also stabilize proteins and membranes. Osmoprotectants play a vital role in helping cells adapt to a variety of harmful environmental situations [25].

The PTS was a major carbohydrate active transport system in bacteria that catalyses the phosphorylation of sugar substrates to cross the microbial cell membrane. 16 genes of TY-11 were related to the PTS **(Additional file 5: Table S4)**. It had Phosphoenolpyruvate-protein phosphotransferase (ptsI encoding PTS system enzyme I) (EC:2.7.3.9). In the first step of PTS system, this enzyme phosphorylates the histidine containing phosphocarrier protein (HPr). Phosphoryl groups from phosphoenol-pyruvate are delivered by HPr to EII enzymes (EIIs), which are also components of TY-11 PTS [33]. Beta-glucosides, cellobiose, fructose, glucitol, galactose, lactose, mannose, sorbitol, and glucose were among the 13 phosphoenolpyruvate dependent PTS EII complexes connected to the transport of carbon sources that were encoded by the genome. The capacity of the TY-11 for transporting carbon is increased by these sugar PTSs’ ability to import several substrates.

#### 7.9. Secretion system

The TY-11 genome featured pieces of the secretion system Sec-SRP. This system includes the signal recognition particle subunit SRP54 (K03106), fused signal recognition particle receptor (K03110) as well as the preprotein translocase subunit SecA (K03070), SecY (K03076), SecG (K03075), and SecE (K03073). Apart from that, the insertion of hydrophobic regions into the lipid bilayer was carried out using membrane protein YidC homologs (LPJCKDFM_00904, LPJCKDFM_01385=K03217). The SecAYEG translocase initially recognizes these membrane proteins. In the process of membrane protein insertion, YidC serves two purposes: in addition to working in concert with the core Sec translocase to aid in protein insertion into the membrane, it can stimulate the insertion of a subset of proteins that do not require the Sec translocase. YidC may aid in the folding of membrane proteins for Sec-dependent proteins throughout the insertion/assembly procedure [34]. Preprotein translocase subunit YajC (K03210) was also available in the secretion system.

#### 7.10. Extracellular matrix, immunity and adhesion

Putative extracellular proteins are one kind of extracellular matrix. In accordance with the existence of a Sec pathway-dependent signal peptide, putative proteins outside the cell of TY-11 were identified (discussed in Section 7.9). Sec proteins serve two purposes: some adhere to cell surfaces and some secrete into the surrounding space. Exopolysaccharides (EPS) are extracellular matrix excreted as tightly attached capsule or loosely bound outer layer slime in microorganisms. It has been proven that exopolysaccharides have a role in the way bacteria communicate with their surroundings. They have been demonstrated to be involved in bacterial biofilm development, adherence to abiotic elements and biotic substrates, and immune system activation [7]. In TY-11 genome, EPS biosynthetic gene clustered from UDP-galactopyranose mutase (K01854), seven enzymes of glycosyltransferases (ko:01003), and one exopolysaccharide biosynthesis protein (LPJCKDFM_00840).

Another point to consider, TY-11 retained a gene for UDP-GlcNAc:undecaprenyl-phosphate/decaprenyl-phosphate GlcNAc-1-phosphate transferase (K02851). N-acetyl-alpha-D-glucosaminyl-diphospho-ditrans, octacis-undecaprenol is a key lipid intermediary for the biosynthesis of numerous microbial cell envelope components (cell wall, cytoplasmic membrane and capsule of gram-positive bacteria), and K02851 catalyzes its synthesis. In some Gram-positive bacteria, the enzyme also starts the manufacture of teichoic acid [35]. Lipoteichoic acids (LTAs) are teichoic acids that are covalently attached to peptidoglycan, whereas wall teichoic acids (WTAs) are teichoic acids that are attached to the lipid membrane [36]. For teichoic acid synthesis (00552 Teichoic acid biosynthesis), the TY-11 harboured 12 genes. Biosynthesis of d-alanyl-lipoteichoic acid requires for enzymatic activation of D-alanin for its incorporation into the membrane associated polymer (mLTA). The TY-11 genome had all five genes of dlt operon for biosynthesis of d-alanyl-lipoteichoic acid: teichoic acid D-Ala incorporation-associated protein DltX (LPJCKDFM_01587), D-alanine—poly (phosphoribitol) ligase subunit DltA (K03367), protein DltB (K03739) involved in the production of D-alanyl-lipoteichoic acid, D-alanine--poly(phosphoribitol) ligase subunit DltC (K14188), and D-alanyl-lipoteichoic acid biosynthesis protein DltD (K03740) **(Additional file 6: Table S5)**. Lipoteichoic Acid (LTA) stimulates nuclear transcription factor kappa B, a component of the innate immune response, after binding to membrane-bound Toll-like receptor 2, according to Nguyen et al. (2017) [37]. In another experiment, the whole genome sequence of *Lactobacillus plantarum* WCFS1 by Grangette et al. (2005) revealed comprehensive bacterial chemicals involved in the interaction with the host immune system. It has been established that peptidoglycan and teichoic acid (TA), particularly lipoteichoic acid (LTA), are involved in the induction of cytokines in the *L. plantarum* cell wall. Toll-like receptor 2-dependent proinflammatory potential of *L. plantarum*-purified LTA was discovered [38]. TY-11 possessed genes for Lipoteichoic Acid (LTA) biosynthesis were predicted for Toll-like receptor 2-dependent innate immune response.

#### 7.11. Flavour related activities

Chemicals that influence the sensations of taste and odour play a major role in determining flavour, which is the sensory perception of food or other things. Glycolysis, proteolysis, and lipolysis are a few of the biochemical procedures used by LAB to create flavor compounds. The flavour-related activities of LAB mostly depend on the species [39]. TY-11 genome possessed pathways of these three types’ metabolisms for producing flavour. For this reason, TY-11 can be used for preparing probiotic yogurt or any other fermented food.

Basically, the sole form of anaerobic homofermentation that the TY-11 genome could perform was homolactic fermentation (details in Section 7.1.3 and Discussion Section), which facilitated the taste of lactic acid, but not acetate and acetaldehyde.

It carried the genes for proteolysis, an important biochemical pathway of flavour production (discussion in Section 7.14-iv). Moreover, transporter proteins for proteolysis, encoded by this genome, were identified as oligopeptide transporter, amino acid branched-chain transport system substrate-binding protein, ion-linked transporter for peptides, and ATP-driven transporter for peptides (discussion in Section 7.8, Additional file 5: Table S4). Bitter peptides are produced during proteolysis, which gives fermented dairy products their bitter flavour. The proteolytically generated amino acids that are free can be transformed into a variety of flavouring substances, such as ammonia, amines, aldehydes, etc [39].

LABs are a source of esterases or lipases, which produces flavour compounds from lipids. TY-11 genome harboured genes for thioesterase (LPJCKDFM_00442), metallophosphoesterase (LPJCKDFM_00966), DHH family phosphoesterase (LPJCKDFM_01373), glycerophosphodiester phosphodiesterase family protein (LPJCKDFM_01556=K01126), phospholipase D-like domain-containing protein LPJCKDFM_01880) for producing flavour compounds from lipids [39].

#### 7.12. Recovering digestive problems within the human intestine

The strain TY-11 with its metabolism capability helped the human gut to digest carbohydrates, lipids, proteins, and other nutrients:

i) Relating with carbohydrate metabolism, its genes for lactose fermentation might be utilized to reduce lactose intolerance (details in Section 7.1.3 and Discussion Section). Lactose intolerance is the major problem globally.
ii) In another point, our observations regarding protein catabolism support its potential role to prevent protein indigestion. Dietary peptides, free amino acids (FAAs), and protein that are ingested enter the large intestine where they are further fermented by the diverse gut flora. The strain TY-11 encoded genes for proteolytic enzymes (discussion in Section 7.14. iv) and peptide transporters (discussion in Section 7.8) to absorb and catalyse small amino acids and peptides. It might affect the host’s energy balance through anaerobic metabolism of peptides and proteins. However, host can also produce some harmful metabolites from proteins in the intestine. This probiotic possessed genes for multicopper oxidase domain-containing protein (LPJCKDFM_00777) and nitroreductase (LPJCKDFM_00111) enzymes to decrease the toxic metabolites (discussion in Section 7.13).
iii) Moreover, this strain was a source of esterases or lipases (section 7.11) and the transmembrane lipid transporter protein (flippases) located in the cell membrane (Section 7.8) to contribute to modulating the human adiposity [40].

Furthermore, the genes, which were responsible for recovering digestive disorders, became apparent in the probiotic TY-11:

i) This is a probiotic that had been shown to help prevent the overgrowth of bad bacteria inside the digestive system, and treat indigestion caused by their overgrowth. The primary mechanism was altering the makeup of gut bacteria through pathogen exclusion and colonization (discussed in Section 7.14).
ii) Similar mechanism might be applicable for alleviating antibiotic-related diarrhoea. Antibiotic-related diarrhoea is caused by disturbing microbial community and their metabolism by antibiotics in the gut. This strain was predicted to relieve antibiotic-related diarrhoea via providing gut barrier by adhering with epithelial cells, preventing harmful bacteria, and replenishing the good bacteria that antibiotics might have killed.
iii) Another digestive issue is irritable bowel syndrome (IBS), which is influenced by changes in flora, enhancing sensitivity to gas in the intestine, increasing inflammation, reducing immune function and changing digestive motility [41].The TY-11 strain obtained the genetic structure for improving IBS symptoms by inhibiting the growth of bad bacteria which produce gas, balancing microbial flora, reducing the gut’s sensitivity to gas build-up by adhering with epithelial cells (discussion in Section 7.14), maintaining acidic pH which controls digestive motility in the gut, and boosting toll-like receptor 2-dependent innate immune response (discussion in Section 7.10) [42]. Kim (2018) described that toll-like receptors play a role to reduce IBS. With decreased acidity, the colon maintains slow peristalsis and motility [43]. The strain TY-11 retained genes to regulate this acidic pH in the gut (discussion in Section 7.14. iv).

#### 7.13. The most prevalent harmful by-products of LABs

It is important to screen for the typical bacterial toxic metabolites that are dangerous to human health and present in LABs. These involve important enzymes, such as D-lactic acid (D-lactate), haemolysins, and biogenic amines (BAs) produced by amino acid decarboxylase (Maria et al., 2008). Although BAs has beneficial impacts on how the body functions, an excessive amount of BAs can be harmful and cause diarrhoea, food poisoning, vomiting, sweating, or tachycardia. Additionally, they can hasten the development of cancer. Histamine (HIS), tyramine (TYR), putrescine (PUT), cadaverine (CAD), tryptamine (TRP), - phenethylamine (PHE), spermidine (SPD), and spermine (SPM) are the most prevalent BAs noticed in cultured foods and drinks. Decarboxylase or deiminase from microorganisms operate to create BAS (Binbin et al., 2020). The TY-11 genome didn’t contain any gene for these metabolites. This is consistent with the biochemical character of *Lactobacillus delbrueckii* ssp. *indicus* WDS-7 [5].

Another aspect, the TY-11 genome carried multicopper oxidase domain-containing protein (LPJCKDFM_00777). Multicopper oxidase (MCO) mediated BAS degradation is an auspicious method as the fermentation process, food nutrition, and flavour are unaffected [44]. This genome also contains nitroreductase (LPJCKDFM_00111), a flavoenzyme that catalyzes the reduction of the nitro groups on nitroaromatic and nitroheterocyclic compounds in a NAD (P)H-dependent manner. For living things, most nitroaromatic chemicals are harmful and mutagenic, however certain microbes, such strain TY-11, have evolved oxidative or reductive mechanisms to break down or modify these substances. Due to its promise uses in bioremediation, biocatalysis, and biomedicine, particularly in prodrug activation for chemotherapeutic cancer therapies, nitroreductases have attracted a lot of attention.

#### 7.14. Characterization of antimicrobial activity from genomic study

The generation of inhibitory chemicals (such as bacteriocin, hydrogen peroxide, and acid), blocking of adhesion sites, competing for resources, and other modes of action are only a few of the ways that probiotics inhibit both Gram negative and Gram positive pathogenic bacteria [45]. The TY-11 genome had genes for following antagonistic mechanism:

i) Three genes commonly are assumed to contribute to H_2_O_2_ production, e.g., pyruvate oxidase, NADH oxidase, and lactate oxidase [46]. Pyruvate oxidase (K00158) was predicted in TY-11, although NADH oxidase, and lactate oxidase were not identified. In enzymology, pyruvate oxidase (EC 1.2.3.3) catalyses the reaction as follows: pyruvate + phosphate + O_2_ ⇌ acetyl phosphate + CO_2_ + H_2_O_2_ This enzyme has pyruvate, phosphate, and oxygen as its three substrates; acetyl phosphate, carbon dioxide, and hydrogen peroxide are its three by-products. The aforementioned enzyme is a member of the oxidoreductase family, especially those that operate on aldehydes or oxo donor molecules with oxygen as the acceptor [47].
ii) The EPS biosynthetic gene clusters of TY-11 help to adhere this probiotic to gut epithelial cells and thereby inhibit the attachment of pathogens with gut by blockage of adhesion sites. EPS biosynthesis was discussed in Section 7.10 and 7.11.
iii) The strain TY-11 can compete with pathogens with its better nutrient (carbohydrate, amino acid, vitamin, minerals, etc.) metabolism and transport capability. These capabilities were discussed in section 7.1, 7.6, 7.7 and 7.8.
iv) Acid released by TY-11 serves in lowering gut pH and preventing the development of pathogenic microbiota. Enzyme genes that regulate post-acidification are (1) genes associated to the metabolism of lactose and genes associated with the pyruvate biosynthesis system (discussion in Section 7.1); (2) the F1F0-ATPase gene cluster (discussion in Section 7.8); (3) genes related to basic amino acid metabolism and synthesis (details in Section 7.6 and Discussion section); (4) genes related to proteins involved in ion transport (discussion in Section 7.8), (5) genes related to enzymes involved in the biosynthetic metabolic pathway of cell membranes (discussion in Section 7.10); and key protease genes in the proteolytic system; among others. A proteolytic enzyme, also known as a protease, proteinase, or peptidase, is any one of the categories of enzymes which breaks down proteins into smaller pieces known as peptides and then into their individual amino acid constituents. The TY-11 uses proteolytic enzymes for generating peptides and releasing free amino acids in order to satisfy its nutritional need [48]. Total thirty genes were responsible for the key proteolysis system and cause TY-11 to have an acidification capability **(Additional file 7: Table S6**).

### 8. Mining bacterial genomic DNA for bacteriocins

To prevent the proliferation of bacterial strains that are identical to or extremely close to them, bacteria create toxins called bacteriocins [49]. Core peptides and their associated proteins were found in the TY-11 genome for synthesizing two bacteriocins. Helveticin-J (331 bp with 46.26% match) of TY-11 showed 99.09% identity and 93% query coverage with helveticin J family class III bacteriocin [*Lactobacillus delbrueckii*] of blast search. Enterolysin_A (275bp with 60.49% match) was also predicted in TY-11 genome (**Fig: 9**).

**Fig: 9.**
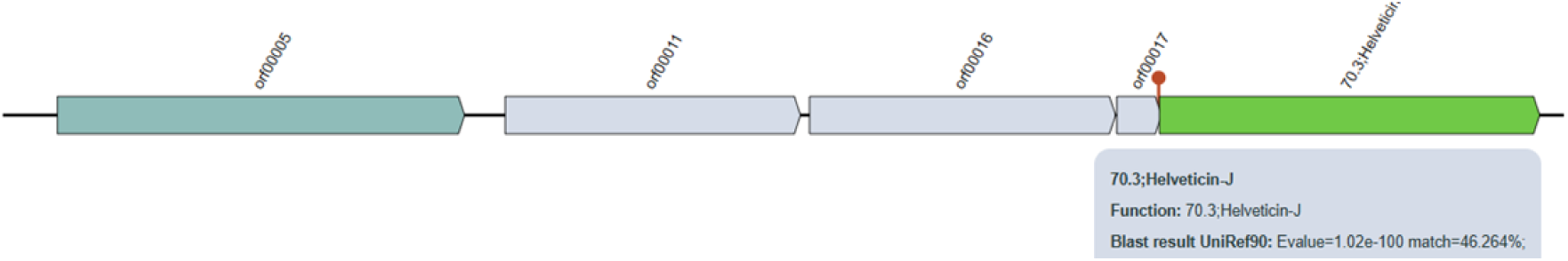
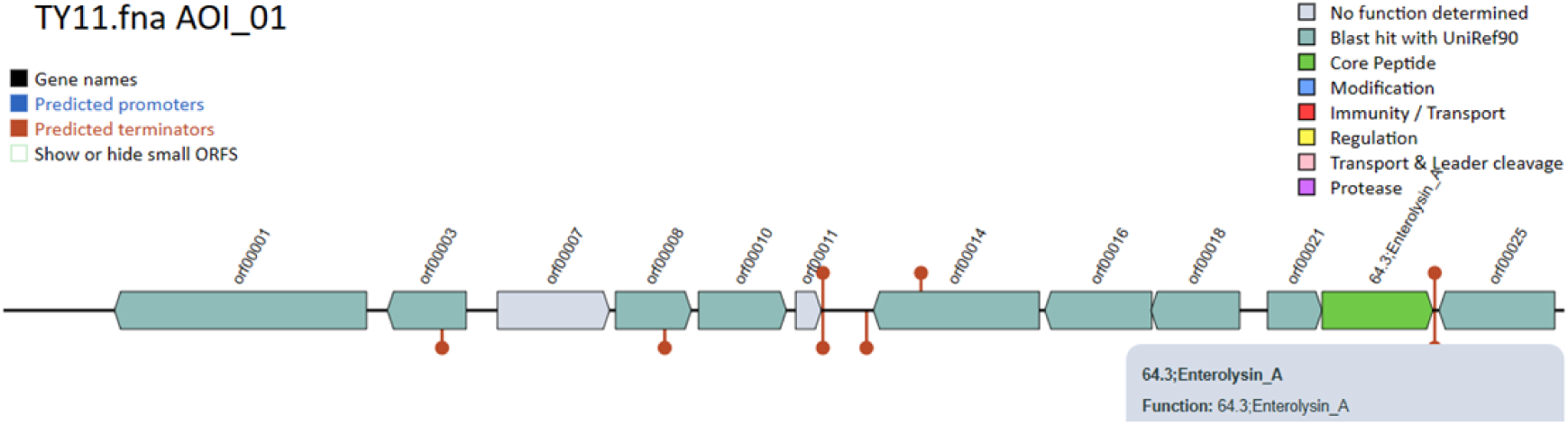
Two bacteriocins in TY-11 genome: Helveticin-J (331 bp), and Enterolysin A (275bp).

### 9. Genomic comparisons

**(TY-11 entire genome represented with *Lactobacillus delbrueckii* core genome, TY-11 non-core genome, and subsp. *indicus* core genome)**

TY-11 entire genome had total 450 annotated known genes. 332 orthologous genes were defined as *Lactobacillus delbrueckii* core genome (263 genomes), and 362 orthologous genes were calculated as subsp. *indicus* core genome (5 genomes). 30 genes were predicted as core genes of subsp. *indicus* (5 genomes) but not core *Lactobacillus delbruickii* genes. These 30 genes are subsp. *indicus* specific genes of TY-11 genome. Remaining 88 genes were TY-11 non-core genome **(Additional file 8: Table S7, Additional file 4: Fig S1)**. The TY-11 non-core genome was further annotated with KEGG database, where 64 genes were annotated (72.7%). Protein families: genetic data processing (8 genes), metabolism of carbohydrates (11 genes), and protein families: signal and cellular functions (12 genes) accounted for the majority of the genes **(Additional file 9: Table S8)**. TY-11 non-core genome had galactose-6-phosphate isomerase subunit LacB (LPJCKDFM_00598), and galactose-6-phosphate isomerase subunit LacA (LPJCKDFM_00599). In enzymology, galactose-6-phosphate isomerase (lacA, lacB; EC5.3.1.26=K01819) is the enzyme which catalyses D-galactose 6-phosphate to D-tagatose 6-phosphate. This enzyme takes role in the metabolism of galactose [50]. Another gene of non-core genome was sucrose phosphorylase encoded by LPJCKDFM_01520-01522. An essential enzyme in the control of various intermediates in metabolic processes and the decomposition of sucrose is known as sucrose phosphorylase (K00690, EC 2.4.1.7). The transformation of sucrose to D-fructose and -D-glucose-1-phosphate is catalysed by it [12]. Although fermentation of sucrose is more variable in subspecies *indicus*, TY-11 bore this gene [4]. The TY-11 non-core genome was composed of 13 genes encoding for CRISPR/CRISPR-associated system (Cas) genes, which were not annotated with KEGG database **(Additional file 8: Table S7**).

## Discussion

The strain TY-11 had been identified as *Lactobacillus delbrueckii* subsp. *indicus*, and its genome (1916674bp) had genes that allowed it to become adapted to the variety of challenging settings found in the human gut. No genes were found for antibiotic resistance or hazardous compounds in this strain.

A family of DNA repeats known as CRISPR offers acquired immunity to foreign genetic components. They generally possess short and extremely conserved repeats (∼30 bp), separated with varying sequences termed spacers, and are very frequently found adjoining to CAS (CRISPR-associated) genes [7]. A total of 45 CRISPR/CRISPR-associated system (Cas) genes of type I, II, and III were detected in the TY-11 genome, which is an unusual occurrence. [51]. Due to current research on the antimicrobial capabilities of type I CRISPR-Cas systems, the TY-11 might be evaluated as a programmable antibacterial in microbiome environments [52].

Twenty genes encoded by phage were predicted in the TY-11 genome. It contained an ATP-dependent chaperone, ClpX that can recognize protein substrates by binding to protein degradation tags. These tags can be short unstructured peptide sequences (e.g., ssrA-tag in *E coli*). As an essential component of ClpP protease complex, ClpX recruits degradable substrates and unfolds their tertiary structure, which requires energy provided by ATP hydrolysis. Subsequently, these ClpX Chaperones transfer protein substrates into the proteolytic chamber formed by ClpP tetradecamer [53]. ClpP proteases contribute to survival and growth of diverse bacteria under conditions of stress [54]. The perfect role of Thymidylate synthase (TS or ThyA) is unidentified within the phage genome but its extra copies probably important for survival of its host by increasing growth [55]. FtsH is not only for the proteolytic removal of unessential proteins, but also in the proteolytic control of some cellular functions [56]. MurA protein catalyses the first step of the peptidoglycan biosynthetic pathway by shifting an enolpyruvate residue to position 3 of UDP-*N*-acetylglucosamine from phosphoenolpyruvate (PEP) [57]. groL is an essential gene (EcoGene: EG10599). The Escherichia coli chaperone protein GroEL (Hsp60) and its regulator GroES (Hsp10) are significant for the proper folding of certain proteins. To function properly, GroEL requires the lid-like cochaperonin protein complex GroES. Both proteins are absolutely essential for bacterial growth at various environmental conditions, such as, heat stress, cold stress, acid stress, and so on [58]. L-lysine is biosynthesized through DAP pathway from DL-2, 6-diaminopimelate. lysA encodes diaminopimelate decarboxylase, which produces L-lysine from meso-diaminopimelate (meso-DAP) [59]. Repressor LexA (EC 3.4.21.88) during transcription represses SOS response genes (encoding principally for cell division inhibitors, DNA repair enzymes and error-prone DNA polymerases) [60]. Through an analogous interaction with the DnaA protein, DnaK participates in the DNA replication of chromosome and hyperosmotic shock [61]. Among the Nus factors, NusA has the most varied functions. It particularly attaches to anti-terminator proteins like N and Q, RNA polymerase (RNAP), nascent RNA, and thus participates in controlling various transcriptional levels: anti-termination, termination and elongation. Also, it takes part in the repairing pathways of DNA [62]. In total, phage proteins have given TY-11 the opportunity to grow and adopt in diverse and stress environments.

The isolate TY-11 harbored genes for glycolysis (Embden-Meyerhof pathway) (M00001), pentose phosphate pathway (M00006) and the three-carbon compound core module of glycolysis (M00002). When glucose is broken down into pyruvate, a little quantity of ATP (energy) and NADH (reducing power) are produced. This process is known as glycolysis. It is a key process that results in the crucial precursor metabolites glucose-6P and fructose-6P, which have six carbons, and glycerone-P, glyceraldehyde-3P, glycerate-3P, phosphoenolpyruvate, and pyruvate, which have three carbons (M00001). The chemical pathways of three-carbon molecules from glycerone-P to pyruvate comprise a conserved core module (M00002) in almost all organisms with fully sequenced genomes; these modules occasionally contain operon structures in bacterial genomes. [63].

It is possible to separate Lactobacilli into two groups: (1) homofermentative species which produce mainly lactic acid (> 65%) from glucose fermentation (e.g. L. acidophilus and L. casei); and (2) species of heterofermentation that generate lactic acid as well as significant amounts of acetate, ethanol and CO2 (e.g. L. fermentum). Through the Embden-Meyerhof-Parnas route, homofermentative LAB preferentially uses hexoses (often glucose). The pentose phosphate pathway may be also present in some homofermentative Lactic Acid Bacteria (LAB) [64]. The pyruvate dehydrogenase and other enzymes required for the transformation of pyruvate into acetaldehyde, acetyl-coenzyme, and acetate were absent from TY-11. It was a homofermentative microorganism that could only produce homolactic fermentation. Homofermentative bacteria are usually used as dairy industry starter cultures. In contrast, heterofermentative bacteria are rarely used as starter cultures within the dairy sector, because they cause defects such as slits in hard cheeses and bloated packaging in other dairy products due to CO2 gas production [65]. Additionally, since the human gut has an anaerobic environment, the absence of pyruvate dehydrogenase complex in strain TY-11 makes sense [6].

Like *Lactobacillus delbrueckii* subsp. *bulgaricus*, TY-11 was able constitutively to degrade lactose but it had no b-galactosidase (EC 3.2.1.23) [3]. This evidence suggests that the strain possessed unique genetic equipment for lactose degradation, which differed from that of the other recognized subspecies. Characterization of this equipment was beyond the scope of the previous studies [4]. lacG gene was found in 5 strains including the strain TY-11 out of 6 strains in subsp. *indicus,* while it was observed in 7 strains out of 69 strains in subsp. *bulguricus* that we used in the phylogenetic tree-1 **(Additional file 4: Fig S1)**. This system is found active in *Lactobacillus delbrueckii* subsp. *lactis.* [66]. Glycoside hydrolase family genes found in this genome (functionally similar to β-galactosidase) have most probably been acquired in the ancestors of *L. delbrueckii* by horizontal gene transfer [30=66]. Worldwide, lactose intolerance in human is a big problem. Treatment of Lactose intolerance in human can include colonic adaptation by prebiotics and probiotics [67]. The TY-11 might be used to lessen the amount of lactose in the intestine of lactose intolerant people.

According to predictions, the TY-11 genome was unable to synthesize the majority of the 20 common amino acids. Some amino acids’ incomplete biosynthesis pathways were present. Although some research suggested that it may also be found in other nutrient-rich settings, such as yogurts [5], this reflects the environment that it can live in the gut, where it may absorb peptides and amino acids from its surrounding environment.

One of the most well-known probiotics, *Lactobacillus delbrueckii* subsp. *bulgaricus* (formerly known as *Lactobacillus bulgaricus*), is the principal bacteria utilized in the manufacturing of cheeses, yogurts, as well as other procedures including naturally fermented goods [68]. In this study, we compared the core genome of subspecies *bularicus* with 69 genomes, and *indicus* with 5 genomes **(Additional file 4: Fig S1)**. 95% threshold was used to compute core genome. We computed 324 and 351 core genes for subspecies *bularicus* and *indicus* respectively. Among these genes only 314 genes were common between subspecies *bularicus* and *indicus* **(Additional file 10: Table S9**).

Only 10 core genes were annotated for “solely *bularicus* but not as *indicus* core genes” **(Additional file 10: Table S9**). Out of 10, 8 genes were present and 2 genes were deficient in the strain TY-11. Phosphate acetyltransferase (pta), and 50S ribosomal protein L36 (rpmJ) were absent in TY-11. Phosphate acetyltransferase (pta) is an enzyme of AckA-Pta pathway. A pathway made up of two enzymes, phosphate acetyltransferase (Pta), which produces acetyl-phosphate from acetyl-coenzyme A, and acetate kinase (AckA), which produces acetate from acetyl-phosphate in a reaction linked to the synthesis of ATP, is responsible for the production of acetate under anaerobic as well as aerobic circumstances. TY-11 lacked Pta, but had AckA in AckA-Pta pathway. Principally, The TY-11 was primarily a strictly homofermentative anaerobic bacteria that could only undergo homolactic fermentation, and did not require maintaining acetate for its metabolism and ATP. It was characteristic for the most lactic acid bacteria, and suited to the gut’s anaerobic environment [6] (discussion in Section 7.1.1). Another protein, ribosomal L36 is the smallest protein of the large subunit (50S) of the bacterial ribosome. It has a specific and key role in the organization of the 23 S rRNA. Corina et al. (2005) speculated that Archaea and Eucarya have used other means to achieve the same degree of stabilization [69]. TY-11 had a dearth of protein L36 and might be compensated with other alternative proteins from 33 large subunit ribosomal proteins encoded in its genome for stabilization and folding of 50S **(Additional file 1: Table S1)**. However, 2 core genes are important which were found in this set of genomes and also in the TY-11 genome. Membrane protein insertase YidC (LPJCKDFM_00904, LPJCKDFM_01385=K03217) encoding gene was one of them. This protein allows the transmembrane segments of the substrate to slide into the lipid bilayer (discussion in Section 7.9). Other core gene of this sort and TY-11 genome was Fe-S cluster assembly protein SufB (LPJCKDFM_01855). This is a fundamental cofactor in proteins engaged in several common cellular functions, including as respiration, central metabolism, ribosome synthesis, DNA replication, and repair [70]. According to our finding, TY-11 genome exhibited similarity with the strains of subspecies *bulgaricus* due to close ANI value. For example, TY-11 complete genome showed 97.24% ANI with the nearest genome sequence of *bularicus* (*Lactobacillus delbrueckii* subsp. *bulgaricus* JCM 1002, GCA_000056065.1).

On the contrary, 37 core genes were predicted only for “at least 95% *indicus* but not for core *bularicus”* **(Additional file 10: Table S9**). TY-11 genome entailed all 37 genes for its function. Complete genome of TY-11 manifested 98.980% ANI with the nearest genome of subspecies *indicus* (*Lactobacillus delbrueckii* subsp. *indicus* JCM 15610, GCA_001908415.1). Because of close ANI value, TY-11 genome demonstrated similarity with the strains of subspecies *indicus*. To take an example, two-component system activity regulator YycH (LPJCKDFM_01012) encoding gene was a crucial gene found in this category. YycH regulates the activity of the essential YycFG two-component system. Cell wall homeostasis is regulated by the YycF-YycG system, suggesting that either an increase or decrease in YycF activity can have an impact on this homeostatic process [71]. Another gene of this group was co-chaperone GroES (LPJCKDFM_00449) (discussion in Section 5). In total, theses 37 genes helped TY-11 and other *indicus* genomes to adapt with environment and to be the exemplary probiotic.

Overall, TY-11 genome contained all the core genes (95% threshold) of subspecies *bularicus* and *indicus,* except two genes: phosphate acetyltransferase (pta), and 50S ribosomal protein L36 (rpmJ). These two genes were not essential for TY-11, and they didn’t extend or constrict any probiotic fingerprint of this genome. The godsend for probiotic idiosyncrasy of TY-11 genome overweighed the blessings of the most promoted subspecies *bularicus* as this had additional 37 genes of subspecies *indicus* core genome.

## Conclusion

According to our knowledge, there are no published articles about functional and comparative genome analysis on *Lactobacillus delbrueckii* subsp. *indicus.* In our current investigation, we employed bioinformatics analysis and comparisons to other probiotic genomes, which revealed a number of common and unique excellences of the probiotic *Lactobacillus delbrueckii* subsp. *indicus* TY-11 that are likely to be important to illustrate its intestinal residence and probiotic roles. Beside genomic studies, phenotypic characters should be unveiled by *in vitro* laboratory tests in further studies.

## Materials and methods

### Isolation and 16S sequencing of bacterial samples

Probiotic species were isolated from a number of samples of yogurts in Tongi, Gazipur, Bangladesh. 1 mL of 10^-6^ decimal diluted sample (suspended in normal 0.9% (w/v) saline solution) was spread on 20-25mL MRS (de Man Rogosa Sharpe, Oxoid, UK) agar medium, and at 37°C the Petri-dishes were incubated for 24-72 h. Based on morphology, several bacterial colonies which showed similarity with *L. delbrueckii* subsp. *indicus* were selected from MRS agar media. In order to purify colonies, the isolates were streaked on the same media and finally the pure colonies were transferred to MRS broth with 15% glycerol for further research. After morphological observation for several samples, only three samples were sorted out for 16S sequencing [72]. Finally, the isolate TY-11 was selected for further experiments.

### Bacterial samples analysis under Transmission Electron Microscope

Samples were applied to pre-hydrophilized carbon-coated EM grids and negatively stained with 1.8% uranyl acetate solution. A VELETA CCD Camera (Olympus Soft Imaging Solutions) placed on a JEM 1010 transmission electron microscope (JEOL) was used to capture images.

### Genome sequencing

Illumina DNA Prep was used to prepare the DNA library of the isolate TY-11. To tagment DNA, this technique, utilizing Bead-Linked Transposomes (BLT), breaks and tags DNA using adapter sequences. The genome of TY-11 was sequenced using a whole-genome (WGS) strategy (30-fold genome coverage) with NextSeq 550 (Illumina, USA).

### QC check of genome sequence

Genome sequence of the isolated TY-11 was checked by FastQC (version 0.11.9), and fastp (version 0.19.7, https://github.com/OpenGene/fastp).

### Genome composition analysis and genome visualization by assembly and annotation

The genome data was assembled using Unicycler 0.5.0 (Wick et al., 2017) and SPAdes 3.15.4 (Prjibelski et al., 2020). Annotation was conducted by PROKKA (version 1.13) [18]. Genome visualization was executed by CGViewBuilder (version 1.1.1) (https://cgview.ca). MobileOG-db was used for searching Mobile Genetic Elements found in bacteria [Brown]. CRISPRCas Finder was used for predicting CRISPR and Cas proteins [73]. The software Resistance Gene Identifier (RGI) (Version 1.1.1) of Comprehensive Antibiotic Resistance Database (CARD) was utilized to predict antibiotic resistance genes [74].

### Phylogenetic study and identification

The new genome was compared with 261 RefSeq genomes of Lactobacillus delbrueckii. All single copy genes that are cotained in the new genome and also present in at least 95% of the total 262 genomes (261 RefSeq genomes + TY-11genome) were selected. These were turned out as 332 genes. The sequences of amino acid of all these genes were extracted and saved into separate files-one file per gene. All amino-acid sequences were aligned in a single file.

Then, nucleotide sequences were re-extracted separately from 332 amino acid sequences. All nucleotide sequences were aligned in a single file. Combined nucleotide sequence alignment for 332 genes from 262 genomes was used for phylogeny. The Neighbor-Joining method was deployed to create a phylogenetic tree [75] implemented in MEGA 11 [76].

As a core genome, 353 genes are present in all 5 *L. delbrueckii* subsp. *indicus* genomes selected from the first tree (TY-11 genome and 4 subsp. *indicus* genomes). The amino-acid sequence alignment was run for those 353 genes (separately for each gene). The Neighbor-Joining approach was employed to infer the second tree employing 5 genomes of *L. delbrueckii* subsp. *indicus* (based on aligned amino-acid sequences of 353 genes) [75] To evaluate tree accuracy, the bootstrap approach of 1,000 bootstrap repeats was utilized. Evolutionary analyses were conducted in MEGA 11 [76].

Identification of the isolated strain TY-11 was conducted by the 16S sequence from the whole genome. The 16S sequence identity was searched by an online search tool, Nucleotide BLAST (https://blast.ncbi.nlm.nih.gov/). We searched Average nucleotide identity (ANI) with complete genome sequence by RAPT (version v0.5.1) (https://www.ncbi.nlm.nih.gov/rapt). Sequence similarity was checked with aligned 353 core gene sequences by Similarity Matrix: MATCH in BioEdit [77].

### Functionality analysis of genome by KEGG annotations

Kyoto Encyclopedia of Genes and Genomes (KEGG) online annotations were used for protein-coding genes of the genome TY-11, using the Blastkola website (https://www.kegg.jp/blastkoala). The PATHWAY database entailed all biological pathway data pertaining to genes. (https://www.kegg.jp/kegg/pathway.html). Each gene’s biological pathway was obtained through KEGG orthology [78].

### Mining bacterial genomic DNA for bacteriocins

Genes and proteins for synthesizing bacteriocins were annotated with the BAGEL database. BAGEL enables researchers to mine bacterial (meta-) genomic DNA for bacteriocins and RiPPs. Additionally, they offer databases as well as a BLAST against the primary peptide databases [79].

Genome comparisons (TY-11 entire genome, *Lactobacillus delbrueckii* core and TY-11 non-core genome, and subsp. *indicus* core genome)

KEGG online annotations were used to compare the functionality of genome compositions.

## Supplementary information

Additional file 1: Table S1. Genes of TY-11 Genome annotated by Prokka.

Additional file 2: Table S2. CRISPR-associated system (Cas) genes.

Additional file 3: Table S3. Phage encoded twenty genes in YT-11 genome.

Additional file 4: Fig S1. Phylogenetic tree-1.

Additional file 5: Table S4. ABC transporters (Ko: 02010).

Additional file 6: Table S5. Extracellular proteins biosynthesis.

Additional file 7: Table S6. Proteolytic enzyme.

Additional file 8: Table S7. 450 annotated orthologous genes of TY-11 genome.

Additional file 9: Table S8. Functional genes of TY-11 non-core genome.

Additional file 10: Table S9. Core genome of subspecies *indicus* and *bulguricus* (combindly).

## Availability of data and materials

The genomes generated and analysed during the current study are available at NCBI under the Project PRJNA975465 at https://www.ncbi.nlm.nih.gov/sra/PRJNA975465. The associated Sequence Read Archive (SRA) and BioSample accession numbers are SRR24696316, and SAMN35326563 respectively.

## Ethics approval and consent to participate

Not Applicable.

## Consent for publication

Not Applicable.

## Acknowledgements

Not applicable.

## Author’s contributions

SMJAM: research proposal, paper writing, lab experiment, raw data and functional genomic analysis, TEM experiment; KK: DNA data analysis, phylogenetic tree, genome sequence annotation, comparative genomic analysis, paper editing; KI: supervision of this project; KS: TEM experiment and analysis, paper review; MEK= plagiarism checking, paper editing, AAIS= plagiarism checking, paper editing, project representative. (Note: Sk. Md. Jakaria Al-Mujahidy=SMJAM, Kirill Kryukov=KK, Kazuho Ikeo=KI, Kei Saito=KS, Md. Ekhlas Uddin=MEK, and Abu Ali Ibn Sina=AAIS).

## References

1. David BA, Nielsen DS, Sørensen KI, Vogensen FK, Sawadogo-Lingani H, Derkx PMF, Jespersen L. *Lactobacillus delbrueckii* subsp. jakobsenii subsp. nov., isolated from dolo wort, an alcoholic fermented beverage in Burkina Faso. Int J Syst Evol Microbiol. 2013; 63(Pt 10):3720–3726. doi: 10.1099/ijs.0.048769-0. Epub 2013 May PMID: 23645015.

2. Kudo Y., Oki K., Watanabe K. *Lactobacillus delbrueckii* subsp. *sunkii* subsp. nov., isolated from sunki, a traditional Japanese pickle. Int J Syst Evol Microbiol. 2012; (62): 2643–2649.

3. Germond JE, Luciane L, Michèle D, Beat M, Giovanna EF, Franco D. Evolution of the Bacterial Species *Lactobacillus delbrueckii*: A Partial Genomic Study with Reflections on Prokaryotic Species Concept. Molecular Biology and Evolution. 2003; 20(1):93–104. 10.1093/molbev/msg012.

4. Dellaglio F, Felis GE, Castioni A, Torriani S, Germond JE. *Lactobacillus delbrueckii* subsp. indicus subsp. nov., isolated from Indian dairy products. Int J Syst Evol Microbiol. 2005; 55(Pt 1):401–404. doi: 10.1099/ijs.0.63067-0. PMID: 15653908.

5. Changjun Wu, Chenwei Dai, Lin Tong, Han Lv And Xiuhong Zhou. Evaluation of the Probiotic Potential of *Lactobacillus delbrueckii* ssp. indicus WDS-7 Isolated from Chinese Traditional Fermented Buffalo Milk *In Vitro*. Polish Journal of Microbiology. 2022; 71(1): 91–105. 10.33073/pjm-2022-012.

6. Liu, Z., Dong, H., Cui, Y. et al. Application of different types of CRISPR/Cas-based systems in bacteria. Microb Cell Fact. 2020; (19):172. 10.1186/s12934-020-01431-z

7. Zhang W, Wang J, Zhang D, Liu H, Wang S, Wang Y and Ji H. Complete Genome Sequencing and Comparative Genome Characterization of *Lactobacillus johnsonii* ZLJ010, a Potential Probiotic With HealthPromoting Properties. Front. Genet. 2019; 10:812. doi: 10.3389/fgene.2019.00812.

8. Brown CL, Mullet J, Hindi F, Stoll JE, Gupta S, Choi M, Keenum I, Vikesland P, Pruden A, Zhang L. mobileOG-db: a Manually Curated Database of Protein Families Mediating the Life Cycle of Bacterial Mobile Genetic Elements. Appl Environ Microbiol. 2022; 88(18):e0099122. doi: 10.1128/aem.00991-22.

9. Van Rossum, T., Ferretti, P., Maistrenko, O.M. et al. Diversity within species: interpreting strains in microbiomes. Nat Rev Microbiol. 2020; 18:491–506 10.1038/s41579-020-0368-1

10. Seemann T, “Prokka: Rapid Prokaryotic Genome Annotation”. Bioinformatics. 2014; 30(14):2068–9.

11. Aisha Farhana; Sarah L. Lappin. Biochemistry, Lactate Dehydrogenase. Treasure Island (FL): StatPearls Publishing. 2022.

12. Elisabeth Coudert, Sebastien Gehant, Edouard de Castro, Monica Pozzato, Delphine Baratin, Teresa Neto, Christian J A Sigrist, Nicole Redaschi, Alan Bridge, The UniProt Consortium. Annotation of biologically relevant ligands in UniProtKB using ChEBI. Bioinformatics. 2023; 39(1). 10.1093/bioinformatics/btac793

13. Eliasson R, Fontecave M, Jörnvall H, Krook M, Pontis E, Reichard P. The anaerobic ribonucleoside triphosphate reductase from Escherichia coli requires S-adenosylmethionine as a cofactor. Proc Natl Acad Sci U S A. 1990; 87(9):3314–8. doi: 10.1073/pnas.87.9.3314.

14. Hao WL, Lee YK. Microflora of the gastrointestinal tract: a review. Methods Mol Biol. 2004; 268:491–502. doi: 10.1385/1-59259-766-1:491.

15. H. Honda, F. Kataoka, S. Nagaoka, Y. Kawai, H. Kitazawa, H. Itoh, K. Kimura, N. Taketomo, Y. Yamazaki, Y. Tateno, T. Saito, β-Galactosidase, phosphor-β-galactosidase and phosphor-β-glucosidase activities in lactobacilli strains isolated from human faeces. Letters in Applied Microbiology. 45(5), 2007; 461– 466, 10.1111/j.1472-765X.2007.02176.x

16. Hengstenberg W, Penberthy WK, Morse ML. “Purification of the staphylococcal 6-phospho-beta-D--galactosidase.” Eur J Biochem.1970; 14(1): 27–32.

17. Lombard, V., Golaconda Ramulu, H., Drula, E., Coutinho, P. M. & Henrissat, B. The carbohydrate-active enzymes database (CAZy) in 2013. Nucleic Acids Res. 2014; 42: D490–D495.

18. Fay A, Glickman MS. An essential nonredundant role for mycobacterial DnaK in native protein folding. PLoS Genet. 2014;10: e1004516. pmid:25058675

19. Susin MF, Baldini RL, Gueiros-Filho F, Gomes SL. GroES/GroEL and DnaK/DnaJ have distinct roles in stress responses and during cell cycle progression in *Caulobacter crescentus*. J Bacteriol. 2006;188: 8044–8053. pmid:16980445

20. Liu J, Huang C, Shin DH, Yokota H, Jancarik J, Kim JS, Adams PD, Kim R, Kim SH. Crystal structure of a heat-inducible transcriptional repressor HrcA from Thermotoga maritima: structural insight into DNA binding and dimerization. J Mol Biol. 2005; 350(5):987–96. doi: 10.1016/j.jmb.2005.04.021. PMID: 15979091.

21. Landsman D. “RNP-1, an RNA-binding motif is conserved in the DNA-binding cold shock domain”. Nucleic Acids Res. 1992; 20 (11): 2861–4. doi:10.1093/nar/20.11.2861

22. McAuliffe, O., R. J. Cano, and T.R. Klaenhammer. Genetic analysis of two bile salt hydrolase activities in *Lactobacillus acidophilus* NCFM. Appl. Environ. Microbiol. 2005; 71:4925–4929.

23. Cotter, P. D., Gahan, C. G. & Hill, C. Analysis of the role of the Listeria monocytogenes F0F1-ATPase operon in the acid tolerance response. Int J Food Microbiol. 2000; 60:137–146.

24. Lorca, G. L., G. Font de Valdez, and A. Ljungh. Characterization of the protein-synthesis dependent adaptive acid tolerance response in *Lactobacillus acidophilus*. J. Mol. Microbiol. Biotechnol. 2002;4: 525–532.

25. Yancey PH. Compatible and counteracting solutes. Strange K Cellular and Molecular Physiology of Cell Volume Regulation. 81109 CRC Press Boca Raton, FL. 1994.

26. Young VR. Adult amino acid requirements: the case for a major revision in current recommendation. J. Nutr. 1994. 124: 1517S–1523S. doi:10.1093/jn/124.suppl_8.1517S.

27. Reeds PJ. Dispensable and indispensable amino acids for humans. J. Nutr. 2000; 130 (7): 1835S–40S. doi:10.1093/jn/130.7.1835S

28. Locasale JW. Serine, glycine and one-carbon units: cancer metabolism in full circle. Nat Rev Cancer. 2013; 13(8):572–83.

29. Yadav MK, Manoli NM, Madhunapantula SV. Comparative Assessment of Vitamin-B12, Folic Acid and Homocysteine Levels in Relation to p53 Expression in Megaloblastic Anemia. PLoS One. 2016; 11(10): e0164559.

30. Aslinia F, Mazza JJ, Yale SH. Megaloblastic anemia and other causes of macrocytosis. Clin Med Res. 2006; 4(3):236–41.

31. Davidson, A. L., Dassa, E., Orelle, C., and Chen, J. Structure, Function, and Evolution of Bacterial ATP-Binding Cassette Systems. Microbiol. Mol. Biol. Rev. 2008; 72, 317–364. doi:10.1128/mmbr.00031-07.

32. Hankins, Hannah M.; Baldridge, Ryan D.; Xu, Peng; Graham, Todd R. Role of Flippases, Scramblases and Transfer Proteins in Phosphatidylserine Subcellular Distribution. Traffic. 2015; 16 (1): 35–47. doi:10.1111/tra.12233

33. Kotrba, P., Inui, M., and Yukawa, H. Bacterial phosphotransferase system (PTS) in carbohydrate uptake and control of carbon metabolism. J. Biosci. Bioeng. 2001; 92, 502–517. doi: 10.1016/S1389-1723(01)80308-X

34. Nil Celebi, Ross E. Dalbey. 4 - YidC: A Protein with Multiple Functions in Bacterial Membrane Biogenesis. The Enzymes. 2007; 25, 93–109. doi:10.1016/S1874-6047(07)25004-8.

35. *Al-Dabbagh B, Mengin-Lecreulx D, Bouhss A.* Purification and characterization of the bacterial UDP-GlcNAc: undecaprenyl-phosphate GlcNAc-1-phosphate transferase WecA. Journal of Bacteriology. 2008; 190 (21): 7141–6. doi:10.1128/jb.00676-08.

36. Swoboda JG, Campbell J, Meredith TC, Walker S. Wall teichoic acid function, biosynthesis, and inhibition. ChemBioChem. 2010; 11 (1): 35–45. doi:10.1002/cbic.200900557.

37. Nguyen, T. T. N., Seo, E., Choi, J., Le, O. T. T., Kim, J. Y., Jou, I., et al. Phosphatidylinositol 4-phosphate 5-kinase α contributes to Toll-like receptor 2-mediated immune responses in microglial cells stimulated with lipoteichoic acid. Cell Signal. 2017; 38, 159–170. doi: 10.1016/j.cellsig.2017.07.009

38. Grangette C, Nutten S, Palumbo E, Morath S, Hermann C, Dewulf J, Pot B, Hartung T, Hols P, Mercenier A. Enhanced anti-inflammatory capacity of a *Lactobacillus plantarum* mutant synthesizing modified teichoic acids. Proc Natl Acad Sci U S A. 2005; 102(29):10321–6. doi: 10.1073/pnas.0504084102.

39. Chen Chen, Shanshan Zhao, Guangfei Hao, Haiyan Yu, Huaixiang Tian & Guozhong Zhao. Role of lactic acid bacteria on the yogurt flavour: A review. International Journal of Food Properties. 2017; 20(1): S316 S330. DOI: 10.1080/10942912.2017.1295988

40. Manasian P, Bustos A-S, Pålsson B, Håkansson A, Peñarrieta JM, Nilsson L and Linares-Pastén JA. First Evidence of Acyl-Hydrolase/Lipase Activity From Human Probiotic Bacteria: *Lactobacillus rhamnosus* GG and *Bifidobacterium longum* NCC 2705. Front. Microbiol. 2020; 11:1534. doi: 10.3389/fmicb.2020.01534

41. Hong SN, Rhee PL. Unraveling the ties between irritable bowel syndrome and intestinal microbiota. World J Gastroenterol. 2014; 20(10):2470–81. doi: 10.3748/wjg.v20.i10.2470. PMID: 24627584; PMCID: PMC3949257.

42. Dai C, Zheng CQ, Jiang M, Ma XY, Jiang LJ. Probiotics and irritable bowel syndrome. World J Gastroenterol. 2013; 19(36):5973-80. doi: 10.3748/wjg.v19.i36.5973. PMID: 24106397; PMCID: PMC3785618.

43. Kim HJ. Do Toll-like Receptors Play a New Role as a Biomarker of Irritable Bowel Syndrome? J Neurogastroenterol Motil. 2018; 24(4):510–511. doi: 10.5056/jnm18153. PMID: 30347933; PMCID: PMC6175550.

44. Ni, X., Chen, J., Du, G. et al. Food-grade expression of multicopper oxidase with improved capability in degrading biogenic amines. Syst Microbiol and Biomanuf. 2022; 2, 285–295. 10.1007/s43393-021-00061-9.

45. Diego Romário Silva, Janaína de Cássia Orlandi Sardi, Nayla de Souza Pitangui, Sindy Magri Roque, Andréa Cristina Barbosa da Silva, Pedro Luiz Rosalen. Probiotics as an alternative antimicrobial therapy: Current reality and future directions. Journal of Functional Foods. 2020; 73(104080). 10.1016/j.jff.2020.104080.

46. Hertzberger, R., Arents, J., Dekker, H. L., Pridmore, R. D., Gysler, C., Kleerebezem, M., & de Mattos, M. J. T. H_2_O_2_ Production in Species of the *Lactobacillus acidophilus* Group: a Central Role for a Novel NADH-Dependent Flavin Reductase. Applied and Environmental Microbiology. 2014; 80(7), 2229–2239. 10.1128/AEM.04272-13

47. Tittmann K, Wille G, Golbik R, Weidner A, Ghisla S, Hubner G. “Radical phosphate transfer mechanism for the thiamin diphosphate- and FAD-dependent pyruvate oxidase from Lactobacillus plantarum Kinetic coupling of intercofactor electron transfer with phosphate transfer to acetyl-thiamin diphosphate via a transient FAD semiquinone/hydroxyethyl-ThDP radical pair”. Biochemistry. 2005; 44 (40): 13291–303. doi:10.1021/bi051058z

48. Cotter, P. D., and C. Hill. Surviving the acid test: Responses of gram-positive bacteria to low pH. Microbiol. Mol. Biol. Rev. 2003; 67:429–453. 10.1128/MMBR.67.3.429-453.2003.

49. Cotter PD, Ross RP, Hill C. Bacteriocins - a viable alternative to antibiotics? Nature Reviews. Microbiology. 2013; 11 (2): 95–105. doi 10.1038/nrmicro2937105.

50. Misselwitz B, Butter M, Verbeke K, Kristin Fox, Mark R. Update on lactose malabsorption and intolerance: pathogenesis, diagnosis and clinical management. Gut. 2019; 68:2080–2091.

51. Horvath P, Barrangou R. CRISPR/Cas, the Immune System of Bacteria and Archaea. Science (New York, NY). 2010;327(5962):167.

52. Gomaa AA, Klumpe HE, Luo ML, Selle K, Barrangou R, Beisel CL. Programmable Removal of Bacterial Strains by Use of Genome-Targeting CRISPR-Cas Systems. mBio. 2014;5(1):e00928–13.

53. Baker TA, Sauer RT. ClpXP, an ATP-powered unfolding and protein-degradation machine. Biochimica et Biophysica Acta (BBA) - Molecular Cell Research. 2012; 1823 (1):15–28.

54. Frees D., Savijoki K., Varmanen P., Ingmer H. Clp ATPases and ClpP proteolytic complexes regulate vital biological processes in low GC, Gram-positive bacteria. Mol. Microbiol. 2007; 63:1285–1295.

55. Lang, Andrew & Hynes, Alexander & Bhattacharjee, Ananda & Woese, Carl & Tariq, Mohammad & Newberry, Fiona & Haagmans, Rik & Booth, Catherine & Wileman, Tom & Hoyles, Lesley & Clokie, Martha & Ebdon, James & Carding, Simon. Genome Characterization of a Novel Wastewater *Bacteroides fragilis* Bacteriophage (vB_BfrS_23) and its Host GB124. Frontiers in Microbiology. 2020; 11. 10.3389/fmicb.2020.583378.

56. Okuno T, Ogura T. FtsH protease-mediated regulation of various cellular functions. Subcell Biochem. 2013; 66:53–69. doi: 10.1007/978-94-007-5940-4_3. PMID: 23479437.

57. Zoeiby El A, Sanschagrin F, Levesque RC. Structure and function of the Mur enzymes: development of novel inhibitors. Molecular Microbiology. 2003; 47:1–12. 10.1046/j.1365-2958.2003.03289.x

58. Keppel F et al. Bacteriophage-encoded cochaperonins can substitute for *Escherichia coli*’s essential GroES protein. EMBO Rep. 2002; 3: 893–8.

59. Dewey, D., Work, E. Diaminopimelic Acid and Lysine: Diaminopimelic AcidDecarboxylase. Nature.1952; 169,533–534. 10.1038/169533a0

60. Erill I, Campoy S, Barbe J. Aeons of distress: an evolutionary perspective on the bacterial SOS response. FEMS Microbiol. 2007; 31 (6): 637–656. doi:10.1111/j.1574-6976.2007.00082.x

61. Calloni G, Chen T, Schermann SM, Chang HC, Genevaux P, Agostini F, Tartaglia GG, Hayer-Hartl M, Hartl FU. DnaK functions as a central hub in the *E. coli* chaperone network. Cell Rep. 2012; 1(3):251–64. doi: 10.1016/j.celrep.2011.12.007. Epub 2012 Mar 8. PMID: 22832197.

62. Sen R, Chalissery J, Muteeb G. Nus Factors of Escherichia coli. EcoSal Plus. 2008; 3(1). doi: 10.1128/ecosalplus.4.5.3.1. PMID: 26443730.

62. Aravind Subramanian, Pablo Tamayo, Vamsi K. Mootha, Sayan Mukherjee, Benjamin L. Ebert, Michael A. Gillette, Amanda Paulovich, Scott L. Pomeroy, Todd R. Golub, Eric S. Lander, and Jill P. Mesirov. Gene set enrichment analysis: A knowledge-based approach for interpreting genome-wide expression profiles. Proceedings of the National Academy of Sciences. 2005; 102 (43): 15545–15550. doi: 10.1073/pnas.0506580102.

63. Gänzle, M.G. Lactic metabolism revisited: metabolism of lactic acid bacteria in food fermentations and food spoilage. Curr Opin Food Sci. 2015; 2, 106–117.

64. Müller, Thomas. Comparison of Methods for Differentiation between Homofermentative and Heterofermentative Lactic Acid Bacteria. ZentralblattFürMikrobiologie, Urban & Fischer. 2012.

65. Hela El Kafsi, Johan Binesse, Valentin Loux et al. Lactobacillus delbrueckii ssp. lactis and ssp. bulgaricus: a chronicle of evolution in action. BMC Genomics. 2014; 15:407. doi:10.1186/1471-2164-15-407

66. Misselwitz B, Butter M, Verbeke K, *Kristin Fox, Mark R.* Update on lactose malabsorption and intolerance: pathogenesis, diagnosis and clinical management. Gut. 2019; 68:2080–2091.

67. Stachelska, Milena Alicja; Foligni, Roberta. Development of a time-effective and highly specific quantitative real-time polymerase chain reaction assay for the identification of *Lactobacillus delbrueckii* subsp. *bulgaricus* and *Streptococcus thermophilus* in artisanal raw cow’s milk cheese. Acta Veterinaria Brno. 2018; 87 (3): 301–308. doi:10.2754/avb201887030301

68. Corina Maeder, David E. Draper. A Small Protein Unique to Bacteria Organizes rRNA Tertiary Structure Over an Extensive Region of the 50S Ribosomal Subunit. Journal of Molecular Biology. 2005; 354(2):436–446. 10.1016/j.jmb.2005.09.072.

69. Blanc B, Gerez C, Ollagnier de Choudens S. Assembly of Fe/S proteins in bacterial systems: Biochemistry of the bacterial ISC system. Biochim Biophys Acta. 2015;1853(6):1436–47. doi: 10.1016/j.bbamcr.2014.12.009.

70. Szurmant H, Nelson K, Kim EJ, Perego M, Hoch JA. YycH regulates the activity of the essential YycFG two-component system in Bacillus subtilis. J Bacteriol. 2005;187(15):5419–26. doi: 10.1128/JB.187.15.5419-5426.2005.

71. Stachelska, Milena Alicja; Foligni, Roberta. “Development of a time-effective and highly specific quantitative real-time polymerase chain reaction assay for the identification of *Lactobacillus delbrueckii* subsp. *bulgaricus* and *Streptococcus thermophilus* in artisanal raw cow’s milk cheese”. Acta Veterinaria Brno. 2018; 87 (3): 301–308. doi:10.2754/avb201887030301

72. Couvin D, Bernheim A, Toffano-Nioche C, Touchon M. et al. CRISPRCasFinder, an update of CRISRFinder, includes a portable version, enhanced performance and integrates search for Cas proteins. Nucleic Acids Res. 2018; 2;46(W1):246-251.

73. Alcock, et al. CARD 2020: Antibiotic Resistome Surveillance with the Comprehensive Antibiotic Resistance Database. Nucleic Acids Research. 2020; 48, D517–D525.

74. Saitou N. and Nei M. The neighbor-joining method: A new method for reconstructing phylogenetic trees. Molecular Biology and Evolution. 1987; 4:406–425.

75. Tamura K, Stecher G, Kumar S. MEGA11: Molecular Evolutionary Genetics Analysis Version 11. Mol Biol Evol. 2021; 38(7):3022–3027. doi: 10.1093/molbev/msab120.

76. Hall, T.A. BioEdit: A User-Friendly Biological Sequence Alignment Editor and Analysis Program for Windows 95/98/NT. Nucleic Acids Symposium Series. 1999; 41:95–98.

77. Kanehisa, M., Sato, Y., and Morishima, K. BlastKOALA and GhostKOALA: KEGG tools for functional characterization of genome and metagenome sequences. J. Mol. Biol. 2016; 428: 726–731.

78. Auke J van Heel, Anne de Jong, Chunxu Song, Jakob H Viel, Jan Kok, Oscar P Kuipers. BAGEL4: a user-friendly web server to thoroughly mine RiPPs and bacteriocins. Nucleic Acids Research. 2018; 46 (W1): 278–281. 10.1093/nar/gky383.

